# Epigenetic signature at *FOXP3* distal enhancer affects regulatory T cell development in Kabuki syndrome

**DOI:** 10.64898/2026.04.08.717184

**Authors:** Alessandra Colamatteo, Antonietta Liotti, Valeria Mazzone, Clorinda Fusco, Antonio Porcellini, Sara Bruzzaniti, Anne Lise Ferrara, Daria Marcogiuseppe, Annamária Szabó, Daniela Melis, Carmelo Piscopo, Matteo Della Monica, Giuliana Giardino, Gioacchino Scarano, Emile Danvin, Bruna De Simone, Francesco Perna, Federica Garziano, Giorgia Teresa Maniscalco, Akshaya Ramachandran, Merve Nida Gokbak, Giuseppe Matarese, Raffaele Iorio, Gilda Varricchi, Giuseppe Spadaro, Giuseppe Merla, Rosa Bacchetta, Irene Cantone, Antonio Pezone, Veronica De Rosa

**Affiliations:** Department of Molecular Medicine and Medical Biotechnology University of Naples “Federico II”, Naples, Italy; Center for Basic and Clinical Immunology Research (CISI) University of Naples “Federico II”, Naples, Italy; Department of Translational Medical Science, University of Naples “Federico II”, Naples, Italy; Institute of Endotypes in Oncology, Metabolism and Immunology (IEOMI) - National Research Council of Italy (IEOMI-CNR), Naples, Italy; “Federico II” University Hospital, 80131 Naples, Italy; Department of Biology University of Naples “Federico II”, Naples, Italy; Department of Medicine Surgery and Dentistry University of Salerno, Salerno, Italy; U.O.S.C. Medical Genetics, A.O.R.N. “A. Cardarelli”, Naples, Italy; Medical and Laboratory Genetics Unit A.O.R.N. San Pio, Benevento, Italy; Department of Agricultural Sciences, University of Naples “Federico II”, Naples, Italy; Department of Advanced Biomedical Sciences, University of Naples “Federico II”, Naples, Italy; U.O.C. Clinical Biochemistry, Ospedali dei Colli, Naples, Italy; Neuroimmunology and Multiple Sclerosis Center “A. Cardarelli”, Naples, Italy; Department of Pediatrics - Stanford School of Medicine, Lorry Lokey Stem Cell Research Building, Office G3039 Stanford, CA, USA; Laboratory of Regulatory and Functional Genomics, Fondazione IRCCS Casa Sollievo della Sofferenza, San Giovanni Rotondo, Foggia, Italy

**Keywords:** DNA methylation, epigenetics, FOXP3, Kabuki syndrome, regulatory T cells, Tregopathy

## Abstract

Kabuki syndrome (KS) is a congenital developmental disorder caused by germinal pathogenic variants in the lysine methyltransferase 2D (KMT2D, KS1) or lysine demethylase 6A (KDM6A, KS2) genes. Kabuki patients display mental retardation, multiorgan malformations and immune dysregulation – ranging from immunodeficiency to autoimmunity – which strongly compromises their life expectancy. We explored whether the complex immunological scenario of Kabuki syndrome 1 subjects (Ks) could be ascribed to an altered generation of CD4^+^FOXP3^+^ regulatory T cells (Tregs). We report that pediatric Ks carrying KMT2D pathogenic variants show a significant reduction of Tregs. DNA methylation analysis reveals a specific methylation pattern at the *FOXP3* distal enhancer that correlates with decreased *FOXP3* transcription early during Treg cell induction and promotes T helper (Th)-2 lineage differentiation. Finally, *in vitro* T cell demethylation rescues FOXP3 expression and Treg induction in Ks, offering a novel potential therapeutic perspective. Our findings connect *KMT2D* loss-of-function to the inhibition of human *FOXP3* gene transcription and provide novel molecular insights to explain the immunological phenotype in Ks, thus pinpointing this syndrome as a novel Tregopathy.

## INTRODUCTION

CD4^+^CD25^+^Forkhead-box-p (FOXP)3^+^ regulatory T cells (Tregs) are essential in the induction and maintenance of immunological homeostasis and *self*-tolerance (Sakaguchi *et al*, 2008). Tregs that develop in the thymus (tTregs) acquire a specific, stable hypomethylation pattern in genes encoding critical molecules, including *FOXP3*, the master transcription factor (TF) that drives Treg development (Lal *et al*, 2009). Alterations of *FOXP3* lead to the development of disorders of the immune response and autoimmune diseases. This is secondary to the inability of the immune system to fight infections due to a disruption of the mechanisms supporting immune homeostasis, which in turn leads to the development of a group of disorders defined primary immunodeficiencies (PID) (Bacchetta *et al*, 2006; Chang *et al*, 2006; Mertowska *et al*, 2022).

Epigenetic modifications of DNA and histones are crucial for the establishment of the chromatin structure in conventional T cells (Tconvs) and direct possible future differentiation into Tregs in the periphery (pTregs) (Kitagawa *et al*, 2013; Lal *et al*., 2009; Placek *et al*, 2017; Zhang *et al*, 2013). *FOXP3* gene transcription is regulated by key distal enhancers - conserved noncoding sequences CNS0, CNS1, CNS2, and CNS3 - that are activated by specific histone modifications, such as acetylation of histone H3 at lysine 27 (H3K27ac) and monomethylation of histone H3 at lysine 4 (H3K4me1) (Bannister & Kouzarides, 2011; Placek *et al*., 2017; Zheng *et al*, 2010). These changes are promoted by a group of enzymes known as the Mixed Lineage Leukemia (MLL) family of histone methyltransferases, which includes MLL1 (KMT2A) and MLL4 (KMT2D)(Allis *et al*, 2007). In humans, loss-of-function pathogenic variants (LoF) of the *KMT2D* gene result in Kabuki syndrome (KS1), a complex and multisystemic developmental disorder (Niikawa *et al*, 1988).

KS1 is characterized by a unique facial dysmorphisms (long palpebral fissures, arched eyebrows, ear anomalies, micrognathia, broad nasal root, and high-arched palate) accompanied by neurological and skeletal abnormalities as well as cardiac, renal, gastrointestinal, endocrine, and immune system alterations (Di Candia *et al*, 2022). KS1 subjects (Ks) frequently meet the “triad” diagnostic criteria of common variable immunodeficiency (CVID), characterized by recurrent airway and gastrointestinal infections, poor response to vaccines and low immunoglobulin levels (Lin *et al*, 2015).

Ks also show up to 54% frequency of autoimmune disorders (Di Candia *et al*., 2022) such as Evans syndrome (immune thrombocytopenia, autoimmune haemolytic anaemia and/or neutropenia), thyroiditis and vitiligo (Ming *et al*, 2005) whose clinical occurrence and treatments are often underestimated (Stagi *et al*, 2016). We explored whether an altered generation of Tregs, the cell subset that finely tunes immune homeostasis (Sakaguchi *et al*., 2008), could be part of the complex immunological picture of Ks and account for the occurrence of autoimmune manifestations in these patients. We examined 12 paediatric Ks – all with *KMT2D* LoF – to assess the phenotype of their blood-derived Tregs. Our findings reveal a significant reduction of Tregs, which correlates with the epigenetic silencing of the *FOXP3* gene, which favors the differentiation toward a Th2-dominant profile. DNA analysis reveals that a specific methylation pattern at the *FOXP3* enhancer region associates with lower FOXP3 expression during the early stages of *in vitro* Treg induction. Our findings highlight a key role for KMT2D in the establishment of the epigenetic landscape necessary for FOXP3 expression, underscoring the importance of monitoring Tregs and immune-dysregulation to enhance clinical management in KS.

## RESULTS & DISCUSSION

To better dissect immune dysregulation in Ks, including abnormal immune profile (Comel *et al*, 2024), we conducted an immunological analysis of the T-cell compartment on 12 pediatric patients, all with *KMT2D* LoF mutations (KS1, Table 1).

**Table 1.** Clinical characteristics of Kabuki subjects. (+) Presence. (-) Absence.

| Kabuki subjects | Patient #1 | Patient #2 | Patient #3 | Patient #4 | Patient #5 | Patient #6 | Patient #7 | Patient #8 | Patient #9 | Patient #10 | Patient #11 | Patient #12 |
| --- | --- | --- | --- | --- | --- | --- | --- | --- | --- | --- | --- | --- |
| Gene | *KMT2D* (MLL4) | *KMT2D* (MLL4) | *KMT2D* (MLL4) | *KMT2D* (MLL4) | *KMT2D* (MLL4) | *KMT2D* (MLL4) | *KMT2D* (MLL4) | *KMT2D* (MLL4) | *KMT2D* (MLL4) | *KMT2D* (MLL4) | *KMT2D* (MLL4) | *KMT2D* (MLL4) |
| Mutation | c.5575delG<br>(p.Asp1859Thrfs*<br>17) | c.14592dupG<br>(p.Pro4865AlafsX<br>48) | c.11714_11716dup<br>AGC<br>(p.Gln3905dup) | c.8196delG<br>(p.Ser2733ValfsX<br>24) | c.705delA<br>(p.Glu237SerfsX<br>24) | c.6595delT<br>(p.Tyr2199IlefsX<br>65) | c.10750C>T<br>(p.Gln3584X) | c.11952dupA<br>(p.Leu3985Thrfs*<br>27) | c.3161_3171delC<br>GTTGAGTCCC<br>(p.Pro1054HisfsX<br>10) | c.16273G>A<br>(p.Glu5425Lys) | r.13593_13671del<br>(p.Leu4532serfs*<br>7) | c.7891G>T<br>(p.Gly2631*) |
| Typical facial feature | + | + | + | + | + | + | + | + | + | - | + | + |
| Autoimmune manifestations | - | Autoimmune<br>Thyroiditis | - | - | - | - | Autoimmune<br>Thyroiditis | - | - | - | IgM anti-cardiolipin | - |
| Allergic/ atopic symptoms | - | - | - | - | + | - | - | - | - | + | - | + |
| Recurrent infections/<br>immune deficiency | + | - | - | IgA deficiency | IgA deficiency | - | - | + | - | + | + | - |
| Skeletal disturbances | Short stature;<br>microcephaly | Short stature;<br>hypodontia; valgus<br>knee | Short stature;<br>microcephaly;<br>scoliosis | - | Hypodontia;<br>scoliosis | Short stature;<br>microcephaly | Scoliosis | - | Malformation of<br>the spine and ribs | Scoliosis | - | Left congenital hip<br>dislocation |
| Mild-to-moderate<br>intellectual/neurological<br>disability | + | + | + | + | + | ++ | + | + | ++ | - | + | + |
| Postnatal growth<br>deficiency | + | + | + | + | - | + | - | - | + | - | - | + |
| Heart disturbances | Aortic coarctation | - | - | Aortic coarctation;<br>interventricular<br>defect; interatrial<br>defect | Anomalous venous<br>return into inferior<br>vena cava; mitral<br>valve prolapse | - | Congenital heart<br>disease | - | Interventricular<br>defect; interatrial<br>defect | Mitral valve<br>prolapse | - | Aortic valve<br>dysplasia |
| Genitourinary<br>diseases | - | Cryptorchidism;<br>hypogonadism | Cryptorchidism;<br>varicocele; severe<br>oligospermia | - | - | - | Enuresis; retractile<br>testes; fused<br>kidneys | - | Cryptorchidism | - | - | Vesical<br>hyperactivity |
| Cleft lip and/or palate | - | - | - | - | - | - | - | - | - | - | - | - |
| Gastrointestinal<br>disturbances | - | - | Inguinal hernia | - | - | Poor feeding | - | - | Poor feeding | - | - | - |
| Hearing and<br>Ophthalmologic<br>disturbances | Bilateral<br>conductive hearing<br>loss; ptosis;<br>strabismus | Bilateral<br>sensorineural<br>hearing loss | - | Mild conductive<br>hearing loss | Bilateral hearing<br>loss | Deafness | Strabismus;<br>hypoaacusis | - | Deafness | Mild conductive<br>hearing loss | - | Ptosis |
| Kidney disturbances | - | - | - | - | - | - | Horseshoe kidney | - | - | - | - | Horseshoe kidney |

When compared to age- and sex-matched healthy subjects (Hs), Ks exhibited a reduction of the CD4^+^ T cell compartment (846±433 in Ks vs. 1097±210 in Hs, Table 2), primarily affecting the naïve CD4^+^CD45RA^+^ T cell subset, both as absolute number (427±338 in Ks vs. 711±218 in Hs, Table 2) and percentage (18.6±8.8 in Ks vs. 25.4±6.2 in Hs, Table EV1), consistently with literature (Comel *et al*., 2024). We did not find any impairment in the CD4^+^ memory compartment in our Ks cohort (Di Candia *et al*., 2022; Lin *et al*., 2015).

**Table 2.** Immunological phenotype of Kabuki subjects (Ks) and healthy subjects (Hs). Absolute numbers per mm^3^ of different immune cell subpopulations in Ks (n=12) and Hs (n=20). Statistical analysis was performed by using Mann-Whitney U-test (two tails) (mean±SD); ns, not significant.

| Cell population (cell/mm<sup>3</sup>) | Ks (n=12) | Hs (n=20) | P |
| --- | --- | --- | --- |
| Leukocytes | 6174±1806 | 6167±1269 | ns |
| Lymphocytes | 2223±1049 | 2837±686 | P=0.0324 |
| CD3<sup>+</sup> | 1631±759 | 2011±469 | P=0.0397 |
| CD4<sup>+</sup> | 846±433 | 1097±210 | P=0.0150 |
| CD8<sup>+</sup> | 610±265 | 682±263 | ns |
| NK<sup>+</sup> | 232±193 | 353±223 | ns |
| B<sup>+</sup> | 309±239 | 432±232 | ns |
| CD4<sup>+</sup>CD8<sup>+</sup> | 31±66 | 28±33 | ns |
| CD3<sup>+</sup>CD45RA<sup>+</sup> | 1071±609 | 1351±445 | ns |
| CD3<sup>+</sup>CD45RO<sup>+</sup> | 560±297 | 660±272 | ns |
| CD4<sup>+</sup>HLA-DR<sup>+</sup> | 16±20 | 17±13 | ns |
| CD4<sup>+</sup>CD45RA<sup>+</sup> | 427±338 | 711±218 | P=0.0090 |
| CD4<sup>+</sup>CD45RO<sup>+</sup> | 419±213 | 386±125 | ns |

Flow cytometry analysis of peripheral blood mononuclear cells (PBMCs) showed that Ks had fewer Tregs relative to Hs, including both canonical CD4^+^FOXP3^+^ Tregs and the more immunosuppressive CD4^+^FOXP3E2^+^ subset, expressing *FOXP3* splicing variants retaining exon2, but comparable FOXP3E2/FOXP3 ratio (Fig. 1a, b; Fig. EV1a-c). Conversely, Ks had a higher frequency of CD4^+^CD25^-^ cells (Fig. 1c). Moreover, CD4^+^FOXP3^+^ Tregs from Ks expressed lower levels of CTLA-4, CD45RA, CCR7, and CD31, but higher levels of PD-1 and CD71, compared to Hs (Fig. 1a and Fig. EV1a). Unlike what previously described, in our cohort of Ks, we observed fewer recent thymic emigrants (CD45RA^+^CD31^+^) and naïve (CD45RA^+^CCR7^+^) cells, yet more CD45RA^-^CCR7^-^cells suggesting an expansion of the effector memory compartment (Fig. 1a and Fig. EV1a) (Barry *et al*, 2022; Di Candia *et al*., 2022; Lin *et al*., 2015). Parallel analysis of CD4^+^FOXP3E2^+^ Tregs showed lower levels of CTLA-4, CD45RA, CCR7, and p-S6, but higher levels of CD71 in Ks (Fig. 1b, Fig. EV1b). This subset also had fewer recent thymic emigrants (CD45RA^+^CD31^+^) and more effector memory (TEMRA) (CD45RA^+^CCR7^-^) cells. The low amount of Treg cells was also confirmed by the low frequency of TSDR demethylated cells in the whole blood of two Ks patients, both with autoimmunity (Patient#2= 0.80%, Patient#7= 0.47% *vs* control average 1.14 ±0.39 % n=52 HS). In addition, also the frequency of the TSDR demethylated cells/CD3 was lower than the normal range (Patient#2 =2.93% and Patient#7= 3.18% *vs* control average 5.45 ±1.79%), therefore showing a decreased ratio between Tregs and Tconvs. Suppressive function of primary sorted CD4^+^CD25^hi^ Tregs was tested in five Ks and only in one of them, who had atopy, we detected a functional defect (Fig. EV1d); however, we cannot exclude that our experimental conditions did not allow to detect small variations as the high 1:1 Treg:Tconv ratio can mask subtle differences in suppressive capacity, as the assay becomes saturated (Akimova *et al*, 2016). Altogether, these data indicate a quantitative defect in Treg cell number, further validated by the low frequency of TSDR-demethylated cells in Ks.

**Fig. 1.**
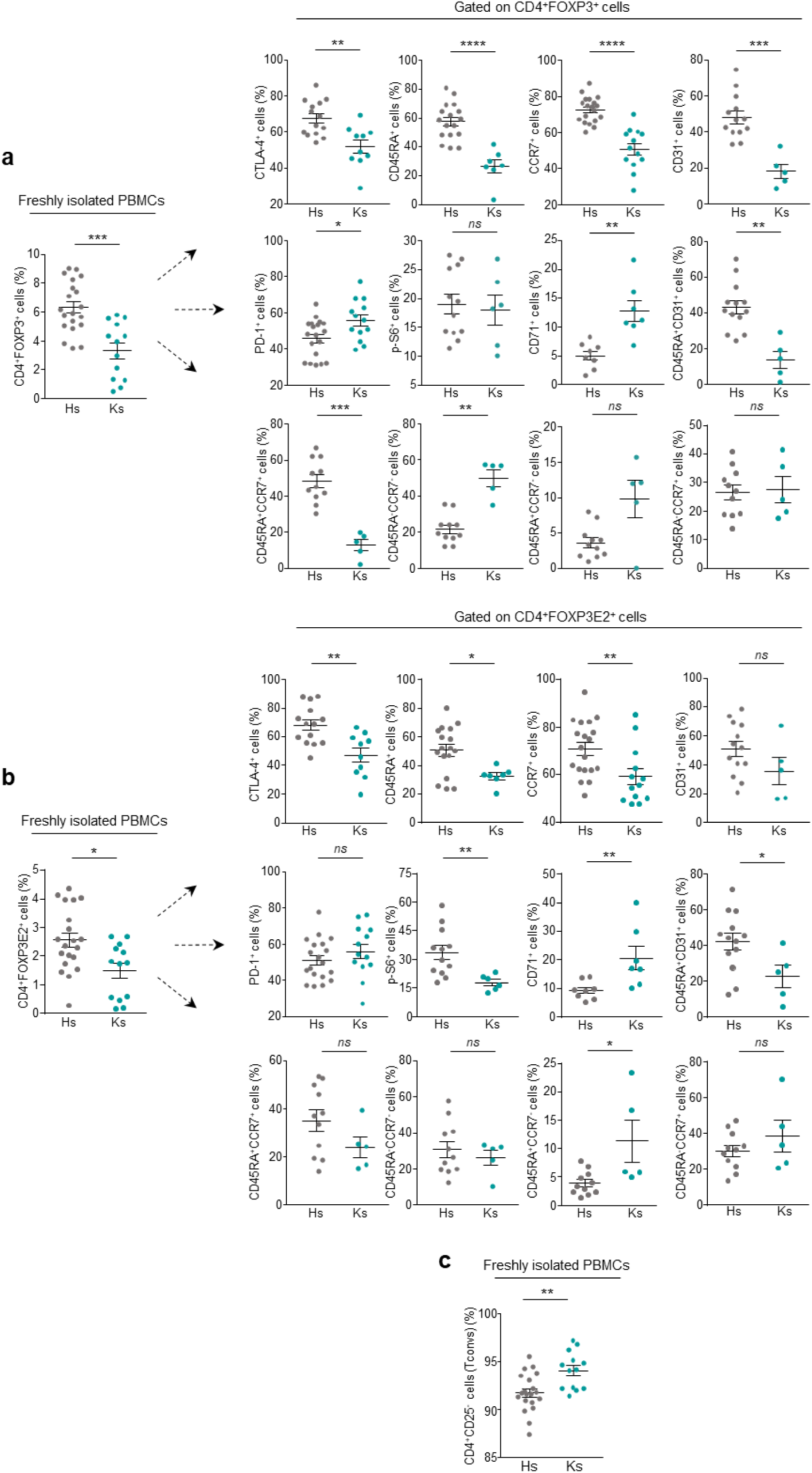
Reduced Treg cell number, altered expression of Treg-cell lineage markers in Ks. Cumulative data showing percentage (%) of peripheral (**a**, upper left panel) CD4^+^FOXP3^+^ Tregs, (**b**, lower left panel) CD4^+^FOXP3E2^+^ Tregs and (**a**,**b**, right panels) expression of Treg-cell specific lineage-markers in freshly isolated PBMCs from Hs (grey dots) and Ks (teal dots) gated on CD4^+^FOXP3^+^ and CD4^+^FOXP3E2^+^ Tregs, respectively. Data are from *n*=20 and *n*=13 independent experiments for Hs and Ks, respectively. Each symbol represents an individual data point (at least 8 Hs and 5 Ks). (**c**) Cumulative data showing percentage (%) of peripheral CD4^+^CD25^-^ cells (Tconvs) in freshly isolated PBMCs from Hs (grey dots) and Ks (teal dots). Data are from *n*=20 and *n*=13 independent experiments for Hs and Ks, respectively. Each symbol represents an individual data point (at least 8 Hs and 5 Ks). Statistical analysis was performed by using Mann-Whitney *U*-test (two tails) (mean±SEM); \**P*≤0.05, \*\**P*≤0.01, \*\*\**P*≤0.001, \*\*\*\**P*≤0.0001; *ns*, not significant.

To explore how KMT2D influences the molecular process underlying the generation of Tregs in Ks, we examined FOXP3 protein expression during the peripheral conversion of Tconvs into inducible (i)Tregs *in vitro* (De Rosa *et al*, 2015). Western blot analysis revealed in Ks a significant reduction of all FOXP3 splicing variants (47- and 44-kDa) and FOXP3E2 isoforms at both time points analyzed (Fig. 2a). Also, their FOXP3E2/FOXP3 ratio was reduced at 24 h in iTregs from Ks (Fig. 2a), thus suggesting that *KMT2D* LoF affects FOXP3E2 expression during Treg cell induction.

**Fig. 2.**
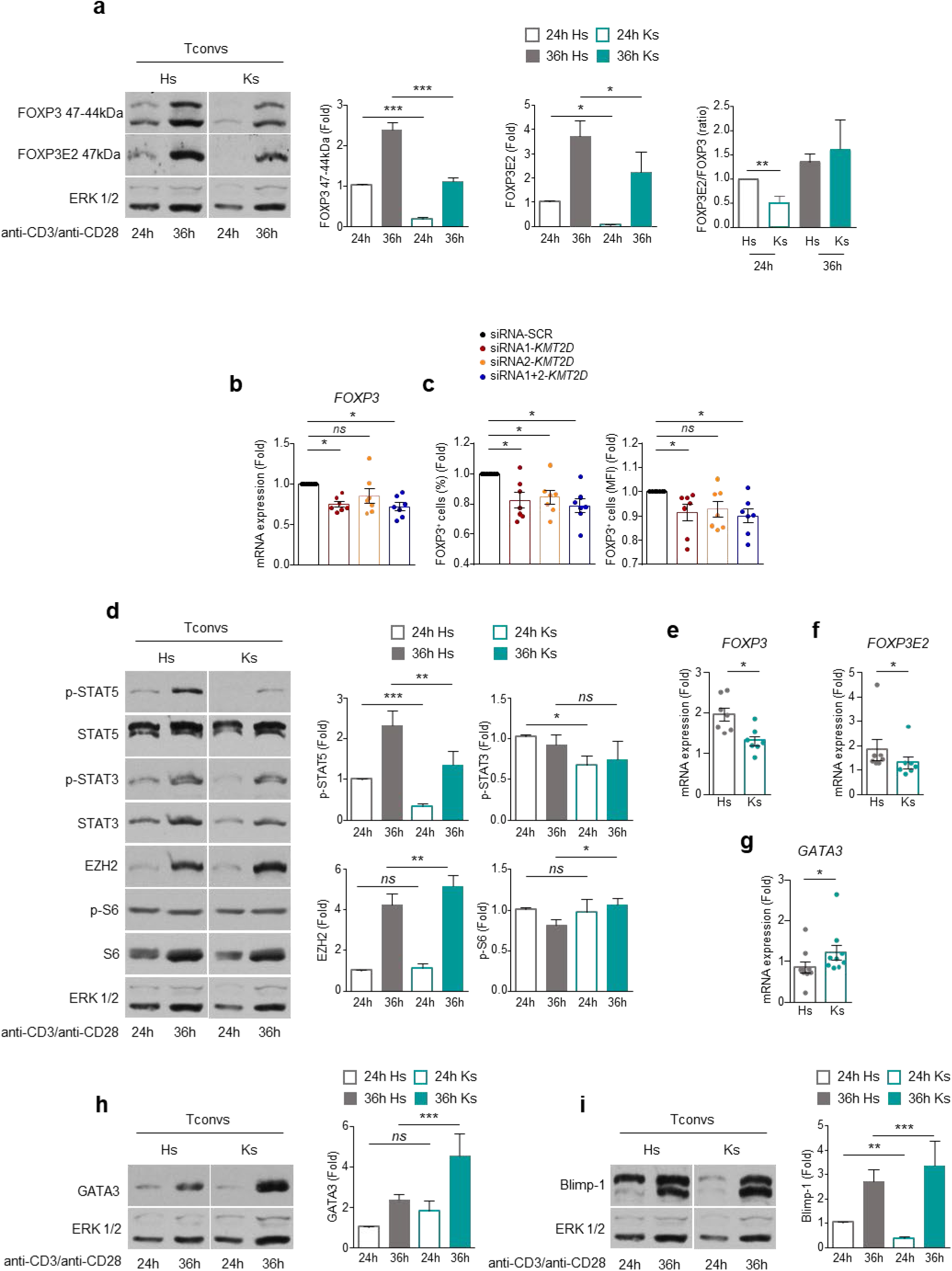
*KMT2D* pathogenic variants affect inducible (i)Treg cell generation and promote Th2 cell differentiation. Western blot analysis of (**a**) FOXP3 (47-44kDa) and FOXP3E2, (**d**) p-STAT5, STAT5, p-STAT3, STAT3, EZH2, p-S6, S6 (**h**) GATA3 and (**i**) Blimp-1 in Tconvs from Hs (empty and full grey column) and Ks (empty and full teal column) after 24h and 36h of TCR stimulation with anti-CD3/anti-CD28 mAbs (0.1 bead per cell). Right, densitometry of (**a**) FOXP3 (47-44kDa), FOXP3E2 and FOXP3E2/FOXP3 ratio (evaluated as the ratio between FOXP3E2 to FOXP3 densitometry values), (**d**) EZH2, (**h**) GATA3 and (**i**) Blimp-1 normalized to total ERK 1/2, (**d**) p-STAT5, p-STAT3 and p-S6 normalized to their total form, presented relative to results obtained for 24h TCR-stimulated Hs-Tconvs (empty grey column). (**a**,**d**,**h**,**i**) Data are from *n*=4 independent experiments from at least 3 Hs and 3 Ks. (**b**) Relative mRNA expression of FOXP3 after silencing of *KMT2D* (siRNA1, siRNA2 and both) in Hs-Tconvs after 24h of TCR stimulation with anti-CD3/anti-CD28 mAbs (0.1 bead per cell). The graph shows *FOXP3* mRNA as fold over siRNA-SCR-Tconvs. (**c**) Cumulative data reporting the percentage (%) (left panel) and mean fluorescence intensity (MFI) (right panel) of FOXP3^+^ cells in CD4^+^CD25^+^ cells after silencing of *KMT2D* (siRNA1, siRNA2 and both) in Hs-Tconvs after 36h of TCR stimulation with anti-CD3/anti-CD28 mAbs (0.1 bead per cell). The graphs are shown as fold over siRNA-SCR-Tconvs. (**b**,**c**) Data are from three independent experiments with technical replicates (*n*=7). Relative mRNA expression of (**e**) *FOXP3*, (**f**) *FOXP3E2* and (**g**) *GATA3* in Tconvs from Hs (grey empty column) and Ks (teal empty column) after 1h of TCR stimulation with anti-CD3/anti-CD28 mAbs (0.2 bead per cell). Data are calculated as fold over unstimulated Hs- and Ks-Tconvs, respectively. (**e**,**f**) Data are from *n*=7 independent experiments from 7 Hs and Ks, respectively. (**g**) Data are from *n*=9 independent experiments from 9 Hs and Ks, respectively. (**e**-**g**) Each symbol represents an individual data point. Statistical analysis was performed by using Wilcoxon test (two tails) (mean±SEM); \**P*≤0.05, \*\**P*≤0.01; \*\*\**P*≤0.005; *ns*, not significant.

KMT2D silencing in Tconvs from Hs, using two different siRNAs, confirmed the direct role of KMT2D in the control of FOXP3 expression during iTreg cell differentiation *in vitro*, both at the mRNA (Fig. 2b) and protein level (percentage and mean fluorescence intensity, MFI) (Fig. 2c).

Treg lineage identity is achieved through epigenetic modifications that control the accessibility of transcription factors (TFs) to regulatory regions of the *FOXP3* gene (Iizuka-Koga *et al*, 2017). Signal Transducer and Activator of Transcription (STAT) and Methyltransferase Enhancer of Zeste Homolog 2 (EZH2) are the main involved factors. STAT promotes CNS2 demethylation *via* Tet methylcytosine dioxygenase (Tet)2 and activates transcription (Ma *et al*, 2018; Pallandre *et al*, 2007; Yue *et al*, 2016), while EZH2 suppresses *FOXP3* by methylating H3K27 and thus reducing chromatin accessibility (DuPage *et al*, 2015). Western blot analysis revealed that Ks have decreased activation of STAT5 and STAT3 and higher EZH2 expression during iTreg induction compared to Hs (Fig. 2d). Additionally, increased p-S6 expression, a downstream target of mTOR known to inhibit FOXP3 expression (Procaccini *et al*, 2010), was also observed in Ks (Fig. 2d). It has been reported that silencing FOXP3 leads to increased expression of genes associated with alternative T cell lineages (Lal *et al*., 2009). We addressed this point and found that FOXP3 impairment during the *in vitro* induction of Tregs from Ks (both FOXP3 and FOXP3E2 mRNAs - Figs. 2e, f), corresponded to an increased transcription of GATA3 - the Th2 lineage marker - at both mRNA (Fig. 2g) and protein level (Fig. 2h). These changes correlated with higher expression of Blimp-1 (Fig. 2i), a zinc finger protein known to promote Th2 differentiation (He *et al*, 2020). Overall, these findings suggest that reduced FOXP3 expression could lead to a preferential induction of Th2 cells in Ks.

KMT2D pathogenic variants markedly decrease mono-methylation of histone H3 lysine 4 (H3K4me) and modify chromatin at the *FOXP3* regulatory regions in mice (Placek *et al*., 2017). Since unmethylated H3K4 can bind and activate the *de novo* DNA methyltransferase (DNMT)3A (Guo *et al*, 2015; Li *et al*, 2011), we investigated whether *KMT2D* pathogenic variants may affect DNA methylation profiles at the regulatory regions of the *FOXP3* gene. We performed targeted deep bisulfite sequencing on three regulatory regions of purified Tconv-DNA: two amplicons for the distal enhancer (CNS0), one for the proximal promoter, and one for CNS2 (Fig. 3a and Table EV2). Our findings revealed no significant differences in average fragment methylation (Fig. EV2a, 3b, 4b, and 6b), Shannon Entropy (Figure EV2b, 3c, 4c, and 6c), single methylated CpG sites (Fig. EV2c, 3d, 4d, and 6d), or taxonomy (Fig. EV2g, 3e, 5a, and 7a) across *FOXP3* regions in Hs and Ks. Notably, qualitative methylation analysis (Pezone *et al*, 2020) of *FOXP3* epialleles (genomic sequences that are identical except for their methylation status) identified families (groups of epialleles sharing a common methylation signature) distinctive for Ks. Cluster analysis of these families (Fig. EV2d, 4e, and 6e) indicated that Hs and Ks epialleles form separate phylogenetic groups (Fig. EV2e). Notably, these epialleles were affected by the position of methylated CpGs (structure), less by their frequency (Fig. EV2f). The proportions of each family within the populations, measured by the MethCore Index (frequency of epiallelic family in the population) and the Clonality Index (frequency of epiallelic family in the methylated population) were comparable between Hs and Ks indicating that they were representative of the population (Fig. 3b, c; Fig. EV4h, i and 6h, i). Structural analysis of these epiallelic families at the distal enhancer (region I) revealed a long, shared cis-methylated core characteristic of the Ks and a shorter, conserved core defining the Hs. These distinct methylated regions, namely Ks-MethCore and Hs-MethCore, represent a specific epigenetic marker of each family (Fig. 3d; Fig. EV4j and 6j). Analysis of the identified epigenetic markers (Ks-MethCore and Hs-MethCore) at the distal enhancer showed that Ks-MethCore frequency was significantly higher in Ks than in Hs (28% *vs* 20%, P<0.05). Conversely, Hs-MethCore was similarly present in both groups (Fig. 3e). Methylation profiling of MethCores at the proximal promoter and CNS2 showed no specific enrichment of epiallelic family in either group (Fig. EV4k and 6k). Due to the high heterogeneity of somatic epialleles (Allen *et al*, 2017; Aref-Eshghi *et al*, 2017; Carter & Zhao, 2021; Landan *et al*, 2012; Morano *et al*, 2014; Pezone *et al*, 2017; Pezone *et al*., 2020; Russo *et al*, 2016), the presence of Ks-MethCore at the *FOXP3* distal enhancer cannot be detected by traditional methods like mean methylation (Fig. EV2a), taxonomy (Fig. EV2g), epiallelic frequencies (Fig. EV2h), or longer epiallele analysis (over 18 CpGs) (Fig. EV8a). Additionally, permutation analysis of Ks-MethCore CpGs supported strong inter-CpG correlation (Fig. EV8b). Overall, the distinct *FOXP3* epiallele family in Ks Tconvs results from *de novo* DNA methylation of specific CpG residues at FOXP3 distal enhancer likely caused by H3K4me depletion in *KMT2D* mutants. Indeed, loss of H3K4me may be compensated by H3K9me/H3K27me, leading to *de novo* DNA methylation of the segment (Placek *et al*., 2017; Russo *et al*., 2016). It is worth noting that *FOXP3* was not included in earlier methylation studies using 450K CpG sites in peripheral blood Ks cells (Aref-Eshghi *et al*., 2017), because no hypermethylated CpGs are present at the *FOXP3* locus in Ks. We propose that the Methylated Core signature at the *FOXP3* distal enhancer suppresses *FOXP3* transcription and affects Treg cell development in Ks. To validate our hypothesis, we treated Ks-Tconvs with azacytidine (aza- a DNA methyltransferase inhibitor) during the generation of iTregs *in vitro*. We found that aza-treatment was able to restore FOXP3 induction in Ks (Fig. 3f), mainly increasing the expression of the FOXP3E2 splicing variants (Fig. 3g).

**Fig. 3.**
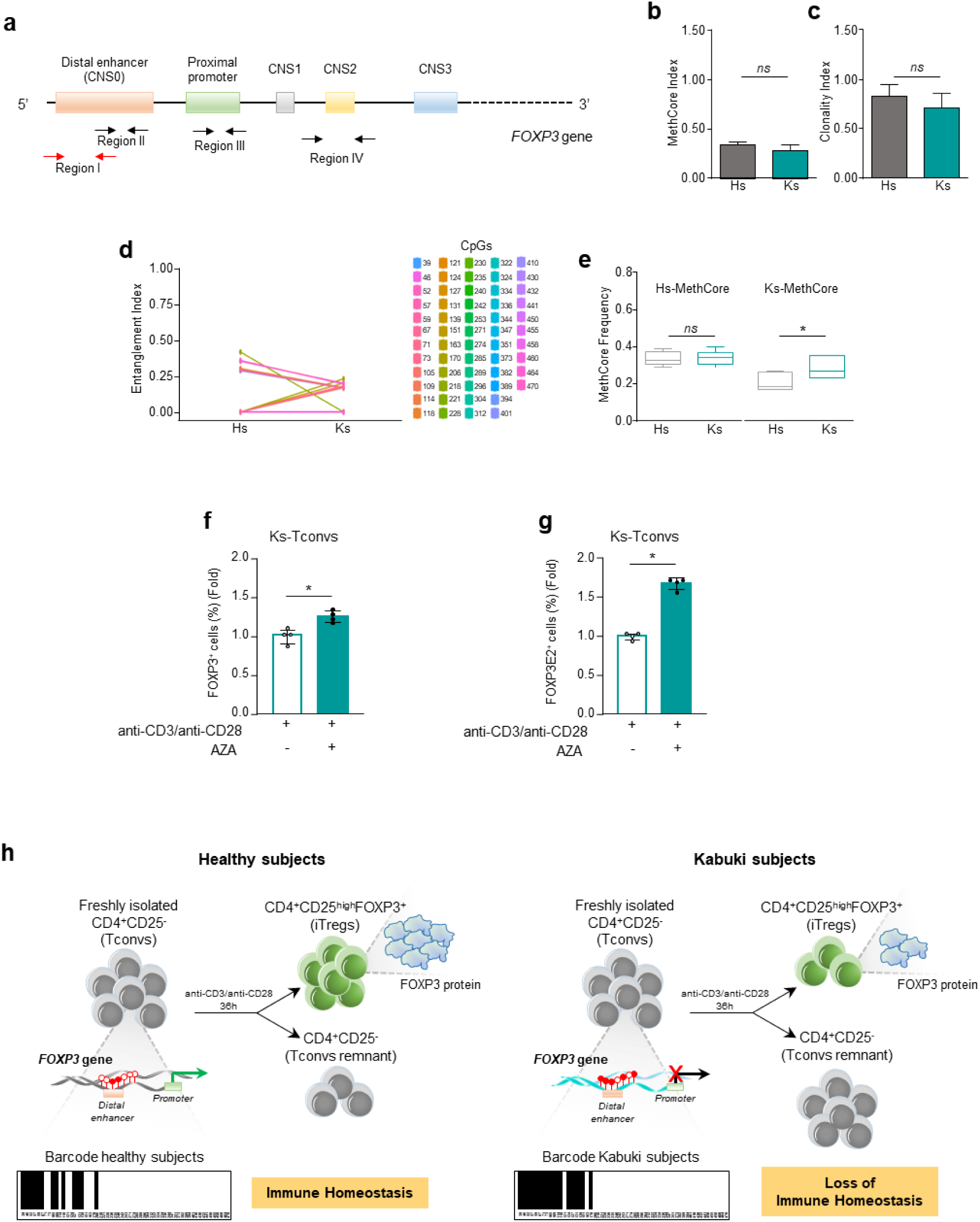
A Methylated Core (MethCore) at the *FOXP3* distal enhancer (CNS0) (region I) marks Tconv cells from Ks. Bisulfite sequencing of DNA extracted from freshly isolated Tconvs from Hs and Ks. (**a**) Human *FOXP3* gene structure: red arrows indicate distal enhancer (CNS0) (region I), black arrows indicate distal enhancer (CNS0) (region II), proximal promoter (region III) and CNS2 (region IV). Analysis of region I refers to the 58 CpGs present. (**b**) Frequency of MethCores (MethCore Index) and (**c**) their normalization to the average methylation (Clonality Index). (**d**) Structure of MethCores of Hs and Ks derived from population of significant molecules. Each CpG is labelled with a color code shown on the right side of the panel (Entanglement Index); the position of CpGs refers to the forward primer. (**e**) Comparison between Hs-MethCore and Ks-MethCore in Hs and Ks, respectively. (**f, g**) FACS analysis determination of FOXP3^+^ and FOXP3E2^+^ Treg generation in Ks-Tconvs stimulated *in vitro* with anti-CD3/anti-CD28 mAbs (0.1 bead per cell) for 4 days, in the presence or in the absence of 5-azacytidine (aza) 5µM. Data are calculated as fold induction over anti-CD3/anti-CD28 stimulated Ks-Tconvs. Data are from two independent experiments with technical replicates (*n*=4) from n=2 Ks. (**h**) Schematic representation of Hs- and Ks-MethCore at the *FOXP3* distal enhancer in Ks. Epigenetic barcode of *FOXP3* distal enhancer, reporting 13 CpGs forming the Hs-MethCore and 18 CpGs forming the Ks-MethCore marked as black lines. 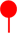 Methylated CpGs. 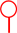 Unmethylated CpGs. Data are from 6 Hs and 6 Ks. Statistical analysis was performed by using the Student’s *t*-test (mean±SD); \**P*<0.05; *ns*, not significant.

In this study, we provide evidence that KMT2D LoF, the genetic cause of KS1, directly affects the epigenetic regulation of *FOXP3* expression, resulting in a low number of Tregs (with low *FOXP3* expression) in Kabuki patients. These results uncover a previously unrecognized mechanism for *FOXP3* gene repression and Treg cell defect in Ks, <u>thus proposing this syndrome as a novel Tregopathy</u> (De Rosa *et al*, 2024). Human *FOXP3* deficiency results in multiorgan failure and autoimmune conditions, leading to IPEX syndrome (immunodysregulation, polyendocrinopathy, enteropathy X-linked). In this disorder, defects in the development, function, or maintenance of Tregs contribute to several autoimmune and inflammatory diseases, due to the loss of immune homeostasis (Alroqi & Chatila, 2016). Their broad regulatory functions are enabled by an array of immunosuppressive mechanisms that regulate both the innate and adaptive immune systems. Our understanding of their pivotal role in immune regulation has critically benefited from the identification of a growing number of inborn errors of immunity (IEIs) known as Tregopathies, which target different pathways governing Treg cell biology and sharing overlapping immune dysregulatory features (De Rosa *et al*., 2024).

Strikingly, Ks often display immune dysregulation during their life, ranging from low immunoglobulin levels and recurrent infections (similar to CVID) (Lin *et al*., 2015) to autoimmune symptoms (predominantly presenting as cytopenia, thyroiditis, and vitiligo), resembling IPEX syndrome (Bacchetta & Roncarolo, 2024; Barzaghi *et al*, 2012) and likely reflecting impaired Treg cell function. Recent studies have linked DNA methylation abnormalities to these clinical features and disturbances in Ks (Sobreira *et al*, 2017), in autoimmune disorders and IPEX patients (Barzaghi *et al*., 2012; Borna *et al*, 2023; Liu *et al*, 2013; Wu *et al*, 2019). Moreover, global methylation profile analysis in Ks revealed a 20% increase in methylation at the promoter of the *homeobox (HOX)-A5* gene, which encodes conserved transcription factors critical for body segmentation and brain patterning during development (Schilling & Knight, 2001). Interestingly, *Kmt2d* knockout models show reduced posterior body length and spinal deformities similar to scoliosis and spina bifida, common in Ks (Van Laarhoven *et al*, 2015). <u>Here, we present evidence that the complex immunological scenario of Ks is caused by an impairment in the differentiation of Tregs, thus suggesting this syndrome as a novel Tregopathy</u>. Indeed, our findings uncover a previously unrecognized mechanism for *FOXP3* gene repression and Treg cell defect in Ks. Using targeted deep bisulfite sequencing, we identified a specific methylation pattern defined “Methylated Core” that suppresses the distal enhancer of *FOXP3* (CNS0) in Tconvs from Ks. This region has also been recently identified as a key IL-2–STAT5 response element that promotes FOXP3 induction in Treg precursors and shows enhancer activity sensitive to methylation-induced silencing (Dikiy *et al*, 2021). The specific combination of epigenetic marks in regulatory regions of lineage-specific genes like *TBX21, GATA3*, *FOXP3*, and *RORγt* influences whether naïve CD4^+^ T cells develop into effector (Teff) or Treg cell subsets (Wei *et al*, 2009). FOXP3 transcription is an early event occurring shortly after TCR stimulation and represents the “primum movens” in T cell lineage decision (Ono, 2020). We observed that repressed *FOXP3* transcription in Tconvs from Ks leads to increased differentiation into GATA3^+^ Th2 cells. A detailed biochemical analysis also revealed lower induction of FOXP3-promoting transcription factors and higher expression of factors supporting Th2 differentiation, such as Blimp-1. The presence of repressive methylation at the *FOXP3* locus could bias Tconv differentiation towards Th2 subsets, which is consistent with the higher incidence of allergic and atopic conditions in Ks (Lin *et al*., 2015) and with other pathologic conditions in which disrupted immune homeostasis associates with higher Th2 frequency (Narula *et al*, 2023; Villa & Notarangelo, 2019).

Consistent with our data, a recent paper demonstrated that in inflammatory conditions, Tregs suppress Th2- as efficiently as Th1- and Th17-responses, even though Th2 responses are the most sensitive to diminished Treg cell number or functionality (Hu *et al*, 2021). Additionally, it has been reported that KMT2D regulates gene expression patterns over space and time through chromatin-based mechanisms and transcriptional compartmentalization (Fasciani *et al*, 2020). Mono-methylation of H3K4 protects against DNA methylation and heterochromatin formation, thereby increasing accessibility and responsiveness to transcriptional activators (Buitrago *et al*, 2021). We found that KMT2D deficiency leads to a DNA methylation pattern that negatively impacts the distal enhancer of *FOXP3*, establishing a new link among the methylated core of the *FOXP3* enhancer, impaired Treg cell development, and immune-related defects in Kabuki syndrome. The impairment is reversible, as we demonstrated that demethylation *via* aza-treatment was able to rescue FOXP3 expression, and this offers new therapeutic perspectives for restoring immune homeostasis in Kabuki syndrome. Our results are further supported by evidence that KMT2D establishes H3K4me1 modification patterns at *FOXP3* regulatory regions, which are essential for its transcription through chromatin looping between distal enhancers and promoters in mice (Placek *et al*., 2017).

It is important to note that, despite the different clinical profile, the functional Treg cell defect was found only in one of the five pediatric patients tested even if this aspect should be confirmed with lower Tconv:Treg ratios (e.g., 1:4, 1:8, or 1:16). Therefore, while the quantitative Treg defect could be an early common phenomenon in KS1, underlying the common immune dysregulation already present in their pediatric stage, the functional defect could occur only later in life, contributing to the onset of autoimmunity. This is in agreement with the increased frequency of autoimmune manifestations observed in adult Ks (Stagi *et al*., 2016). In line with this, the different grade of impairment observed by MethCore profile could hide a pathogenic condition that they will develop in adult life.

In summary, our findings indicate that mono-methylation by KMT2D is crucial for CpG demethylation at the *FOXP3* upstream enhancer, preparing it for transcriptional activation when conditions are favourable in naïve CD4^+^ T cells (Fig. 3h). These results offer new insights into the molecular basis of Ks and suggest that epigenetic and DNA methylation patterns may serve as targets for developing new treatments for immune dysregulations in Kabuki syndrome.

## Methods

### Subjects and study design

Clinical and demographic characteristics of our study cohort are shown in Table 1 and Table EV3, respectively. Human subjects enrolled in this study were collected after obtaining informed consent. The study was approved by the Institutional Review Board of University of Naples “Federico II” (Protocol n. 304/16). Kabuki and healthy subjects (Ks and Hs) were enrolled by clinicians at U.O.S.C. Medical Genetics, A.O.R.N. “A. Cardarelli”, Naples; Medical Genetics Unit A.O.R.N. San Pio, Benevento and Department of Translational Medical Science, Section of Pediatrics, University of Naples “Federico II”, Naples. Ks were diagnosed by clinical manifestations and confirmed molecularly by genetic *KMT2D* mutation screening (all with *KMT2D* loss-of-function). Blood samples were processed within 4 hours.

### Immunophenotypic analysis

Blood samples from Ks and Hs were collected in heparin-tubes and were used for immune cell profiling (Table 2 and Table EV1). Whole blood cells were analysed with a clinical-grade haematocytometer to determine absolute lymphocyte numbers in each sample. One-hundred μl of blood was incubated 30 min at room temperature with the specific antibody combinations. Red blood cells were lysed using IOTest 3 Solution (Beckman Coulter Life Sciences) for 15 min and samples were subsequently washed and resuspended in 300 μl phosphate-buffered saline (PBS). Flow cytometry was carried out by using AQUIOS Tetra-1+ Panel Monoclonal Antibody Reagents consisting of CD45-FITC (B3821F4A)/CD4-RD1 (SFCI12T4D11)/CD8-ECD (SFCI21Thy2D3)/CD3-PC5 (UCHT1) and AQUIOS Tetra-2+ Panel Monoclonal Antibody Reagents consisting of CD45-FITC (B3821F4A)/(CD56 + CD16)-RD1 (3G8+N901)/CD19-ECD (J3-119)/CD3-PC5 (UCHT1), PC7-anti-HLA-DR (Immu-357) and ECD-anti-CD45RO (UCHL1) (all from Beckman Coulter Life Sciences). Flow cytometry was carried out on cells gated on CD45^+^ - Side Scatter (SSC). Immunophenotypic analysis was performed with Cytomics FC500 Flow Cytometer (Beckman Coulter Life Sciences) with CXP software (Version 2.0) and Kaluza Analysis Flow Cytometry software (Version 2.1.1).

### Cell purification, cultures and generation of inducible (i)Treg cells

Peripheral blood mononuclear cells (PBMCs) from Ks and Hs were isolated after Ficoll-Hypaque gradient centrifugation (GE Healthcare). Peripheral Treg (CD4^+^CD25^+^CD127^-^) and Tconv (CD4^+^CD25^-^) cells were purified (90-95% pure) from PBMCs of Hs and Ks by magnetic cell separation with a Regulatory CD4^+^CD25^+^ T Cell Kit (Invitrogen). For suppression assays, Tregs and Tconvs were cultured (1 x 10^4^ cells per well) for 60h in round-bottom 96-well plates (Corning Falcon) in RPMI 1640 medium (Gibco) supplemented with penicillin (100 UI/ml), streptomycin (100 µg/ml) (Gibco) and 5% autologous serum (AS) and stimulated with Dynabeads coated with mAb to CD3 plus mAb to CD28 (anti-CD3/anti-CD28 mAbs) (0.5 bead per cell; Gibco). After 48h, Thymidine, (methyl-^3^H) (0.5 μCi/well; Perkin-Elmer) was added to the cell cultures, and cells were harvested 12h later. Radioactivity was measured with a β cell plate scintillation counter (Wallac). Percentage suppression of Tconv cell proliferation was calculated as: (cpm of Tconvs alone – cpm of Tconvs treated with Tregs)/cpm of Tconvs alone) *100, as already reported(Collison & Vignali, 2011). For the generation of inducible (i)Tregs, Hs- and Ks-Tconvs and *KMT2D*-silenced Hs-Tconvs were cultured (2 x 10^5^ cells per well) in round-bottom 96-well plates (Corning Falcon) with RPMI-1640 medium (Gibco) supplemented with penicillin (100 UI/ml), streptomycin (100 µg/ml) (Gibco) and 5% AS or 5% AB human serum (EuroClone) and 1h-stimulated (0.2 bead per cell; Thermo-Fisher Scientific) or 24h-and 36h-stimulated (0.1 bead per cell; Thermo-Fisher Scientific) with anti-CD3/anti-CD28 mAbs. For 5-azacytidine (aza) experiments, Ks-Tconvs were cultured (1 × 10^6^ cells per ml) in 96-well plates (Corning Falcon) with RPMI-1640 medium (Gibco) supplemented with 100 UI/ml penicillin, 100 μg/ml streptomycin (Gibco) and 5% AS serum at a density of 0.1 bead per cell for 4 days, in the presence or in the absence of 5 µM aza. Tconvs were then harvested and used for FACS analyses after staining with the following mAbs: APC-H7–conjugated anti-human CD4 (RPA-T4), phycoerythrin (PE)–indodicarbocyanine (Cy5)–conjugated anti-human CD25 (M-A251), PE-conjugated anti-human FOXP3 from eBioscience (PCH101, that recognizes all splicing variants through an epitope of the amino terminus of FOXP3) and PE-conjugated anti-human FOXP3 from eBioscience (150D/E4, that recognizes FOXP3E2 variants through an epitope present in the Exon 2 only).

### Flow cytometry

Freshly isolated PBMCs from Ks and Hs or 36h-stimulated (anti-CD3/anti-CD28 mAbs 0.1 beads per cell) *KMT2D*-silenced Hs-Tconvs were surface stained with the following mAbs: APC-H7–conjugated anti-human CD4 (RPA-T4), PE-Cy7–conjugated anti-human CD25 (M-A251), FITC–conjugated anti-human CD71 (M-A712), PerCP-Cy5.5–conjugated anti-human CD197/CCR7 (150503), BB515–conjugated anti-human CD45RA (HI100), BV421–conjugated anti-human CD279/PD-1 (EH12.1), BV510–conjugated anti-human CD31 (WM59) (all from BD Biosciences). Thereafter, cells were washed, fixed, and permeabilized (Human FOXP3 Buffer Set; BD Biosciences) and stained with following mAbs: PE–conjugated anti-human FOXP3 (259D/C7), APC–conjugated anti-human CD152/CTLA-4 (BNI3) (all from BD Biosciences), PE–conjugated anti-FOXP3 (150D/E4) (from Invitrogen) and Alexa Fluor 647–conjugated antibody to ribosomal protein S6 phosphorylated at Ser235 and Ser236 (D57.2.2E) (from Cell Signaling Technology). Flow cytometry data were analysed with FACSCanto II (BD Biosciences) with DIVA software (Version 6.1.3) and FlowJo software (Tree Star, Version V10).

### Molecular signaling and immunoblot analysis

To generate iTreg cells, Ks- and Hs-Tconvs were stimulated with anti-CD3/anti-CD28 mAbs (0.1 beads per cell) for 24h and 36h. Total cell lysates were obtained incubating cells for 20 min at 4°C in RIPA assay buffer (Sigma-Aldrich), plus SIGMAFAST Protease Inhibitor (Sigma-Aldrich) and Sigma Phosphatase Inhibitor (P5726; Sigma-Aldrich), and immunoblot analyses were performed as described(De Rosa *et al*, 2007). The mAbs used were the following: anti–phospho-STAT5 (Tyr694) and anti-STAT5 (D206Y), anti–phospho-STAT3 (3E2) (Tyr705) and anti-STAT3 (79D7), anti-EZH2 (D2C9), anti-GATA-3 (D13C9), anti-Blimp-1 (C14A4), anti-S6 phosphorylated at Ser240 and Ser244 and anti-S6 (54D2) (1:1000 dilution; all from Cell Signaling Technology); anti-FOXP3 (PCH101) and anti-FOXP3E2 (150D/E4) (1:500 dilution; all from Invitrogen); and anti-ERK 1/2 (H72) (1:1000 dilution; Santa Cruz Biotechnology) to normalize the amount of loaded protein. All filters were quantified by densitometry using ImageJ software (NIH). We scanned at least two films with different exposures for each protein detected. Results were calculated as the densitometry of phosphorylated protein normalized to its total form (for STAT5, STAT3 and S6) or normalized to densitometry of ERK 1/2 (for FOXP3, FOXP3E2, EZH2, GATA3 and Blimp-1) and are presented relative to results obtained for 24h TCR–stimulated Tconvs from Hs.

### TSDR analysis using epigenetic quantitative PCR testing protocol

Peripheral whole blood or dried blood spots (DBS) were lysed at 56 °C for 15 minutes. DNA isolation and bisulfite conversion were performed using BisuPrep Reagents (RUO; Epimune Diagnostics). The bisulfite-converted DNA was similarly isolated from sorted FACS-sorted cell pellets. The eluted DNA was then used to perform quantitative PCR using specific markers for FOXP3^+^ Tregs, CD3^+^ T cells, CD4^+^ T cells, glyceraldehyde-3-phosphate dehydrogenase, and GAP [GC] controls, and the quantification of cell subsets was performed as already described (Baron *et al*, 2018; Borna *et al*., 2023).

### Quantitative real-time PCR

Total cellular RNA was extracted from Ks- and Hs-Tconvs freshly isolated, 1h-stimulated anti-CD3/anti-CD28 mAbs (0.2 bead per cell) or from Hs *KMT2D*-silenced Tconvs 24h-stimulated with anti-CD3/anti-CD28 mAbs (0.1 bead per cell), by using Arcturus^TM^ PicoPure^TM^ RNA Isolation Kit (Applied Biosystem). cDNA was synthetized in a 20 µl reaction volume containing 0.5 µg of total RNA and SuperScript IV VILO (Invitrogen). *FOXP3* all transcripts, *FOXP3* splicing variants containing the Exon 2 (*FOXP3E2*), *GATA3* and *KMT2D* mRNAs were detected by using TaqMan Universal Master Mix II, with UNG (Applied Biosystem) according to the manufacturer’s instruction, and analysed with QuantStudio3 Real-Time PCR System (Applied Biosystem). As internal standard control to perform normalization between samples we used *18S* ribosomal RNA. TaqMan probes were from Applied Biosystem (Table EV4). The mRNA expression was expressed as relative amount compared to *18S* RNA using the ΔCt method (2^-ΔCt); ΔCt is the difference in Ct between the gene of interest (*FOXP3* all transcripts, *FOXP3E2*, *GATA3* and *KMT2D*) and the endogenous control (*18S*). Results were calculated as fold over freshly isolated Tconvs from Ks and Hs or as fold over siRNA-scramble (SCR)-Tconvs from Hs, respectively.

### DNA extraction, bisulfite conversion, PCR and sequencing

Genomic DNA was extracted from freshly isolated Ks- and Hs-Tconvs by using PureLink™ Genomic DNA Mini Kit (Invitrogen). Bisulfite treatment of genomic DNA was performed by using EZ DNA Methylation-GoldTM Kit (Zymo Research). PCR was performed in a final volume of 20 μl containing: 1x reaction buffer, 0.5 U FastStart High Fidelity Taq polymerase (Roche), 0.2 mM dNTP mix (Roche), 0.2 μM for each forward and reverse primers(Kennedy *et al*, 2014) (Table EV5), and 2 μl of bisulfite-treated genomic DNA under the following thermo cycle conditions: one cycle at 95°C for 5 min followed by 5 cycles at 95°C for 45 sec, 5 cycles at 55°C for 40 sec, 5 cycles at 72°C for 45 sec, followed by 35 cycles at 95°C for 45 sec, 35 cycles at 52°C for 40 sec, 35 cycles at 72°C for 45 sec and a final extension step of 10 min at 72°C (region I and region IV); one cycle at 95°C for 5 min followed by 5 cycles at 95°C for 45 sec, 5 cycles at 56°C for 40 sec, 5 cycles at 72°C for 45 sec, followed by 35 cycles at 95°C for 45 sec, 35 cycles at 53°C for 40 sec, 35 cycles at 72°C for 45 sec and a final extension step of 10 min at 72°C (region II); one cycle at 95°C for 5 min followed by 5 cycles at 95°C for 45 sec, 5 cycles at 54°C for 40 sec, 5 cycles at 72°C for 45 sec, followed by 35 cycles at 95°C for 45 sec, 35 cycles at 50°C for 40 sec, 35 cycles at 72°C for 45 sec and a final extension step of 10 min at 72°C (region III). Three μl of each PCR products were used to check product size on 1.2% agarose gel. PCR products were purified by using QIAquick PCR Purification Kit (QIAGEN) and were pooled at equimolar ratio. DNA sequencing was performed by IGA technology (IGATech) services (Udine, Italy), using Illumina MiSeq sequencing.

### *KMT2D* silencing

For *KMT2D* knockdown, Hs-Tconvs were transiently transfected with a Neon Transfection System (Invitrogen) with siRNAs designed to target two different regions of *KMT2D*: siRNA1 (ID: s15604; 4392420; Invitrogen) or siRNA2 (ID: s15606; 4392420; Invitrogen) or with scrambled sequence (4390843; Invitrogen) at final concentration of 200 nM (in medium without serum). Tconvs were transfected under the following conditions: pulse voltage, 2,200 V; pulse width, ms; pulse number, 1. Then, cells were stimulated with anti-CD3/anti-CD28 mAbs (0.1 beads per cell) for 36h. After 24h of stimulation, quantitative real-time PCR was performed for confirmation of specific silencing (Fig. EV9).

### Bisulfite sequencing analysis

Paired sequences in FASTQ format by Illumina sequencing machine were filtered and assembled using Paired-End reAd mergeR (PEAR) (Zhang *et al*, 2014). FASTA format were obtained using PReprocessing and INformation of SEQuence (Prinseq) (Schmieder & Edwards, 2011). Reads were aligned to the bisulfite converted reference sequence. Reads with ambiguous calls at the CpG dinucleotide were removed. To extract mCpG configurations in single DNA molecules, reads in FASTA format were processed using ampliMethProfiler (Camacho *et al*, 2009; Caporaso *et al*, 2010; Scala *et al*, 2016) applying several quality filters. In particular, we retained only reads characterized by (i) length ± 50% compared with the reference length, (ii) at least 80% sequence similarity of the primer with the corresponding gene, (iii) at least 98% bisulfite efficiency and (iv) alignment of at least 60% of their bases with the reference sequences. The methylation status of all cytosines in the CpG sequence context is coded as methylated (1) or unmethylated (0). Reads with ambiguous calls (including gaps or A or G) at the CpG dinucleotide were removed. The data, in binary formats, were successively analysed with the MethCoresProfiler (available at https://github.com/84AP/MethCoresProfiler/) (Pezone *et al*., 2020). Briefly, MethCoresProfiler extracted epialleles containing combinations of 2-CpGs whose observed frequencies (population count) are significantly higher or lower than expected (product of independent events) by applying a chi-square. It also removed all epialleles whose frequencies were equal to a random sample created with the same parameters as the sample under examination (number of CpGs and sample size). Then, it extracted the maximum combination of CpGs shared among all selected epialleles and performed a cluster analysis for frequency and phylogeny. It also calculated three indices: (i) frequency of the core in the population, (ii) frequency of the core with respect to the average of methylation and (iii) frequency of the CpGs constituting the core with respect to the average methylation of the individual CpGs. The *P* value used by MethCoresProfiler is 10^-10.

### Stress Test Methodology

To confirm the goodness of the Ks-MethCore, we performed three tests. (i) Search for random combinations formed by 18 CpGs structurally different from the Ks-MethCore (Random18) did not show significant differences within the two populations (Hs and Ks). (ii) Frequencies of methylated classes with more than 18 CpGs at the *FOXP3* distal enhancer did not show significant differences within the two populations (Hs and Ks) (Fig. EV8a). (iii) MethCoresProfiler identified a MethCore of 18 CpGs in Ks from a population with 258 possible combinations of epialleles. To evaluate the strength of the association of CpGs within the Ks-MethCore, we performed “One-point permutation test” by adding or removing a single CpG in the Ks-MethCore. Among all the 739 possible combinations obtained with the 58 CpGs present in the region I, only 151 were significantly more frequent in Ks than in Hs (Fig. EV8b). Analysis of the CpGs making up the 151 combinations shows that the 18 CpGs of the Ks-MethCore are present with a frequency of 0.95; *P* value < 0.05. No other CpGs external to the Ks-MethCore show such a strong association (Fig. EV8b).

### Statistical analysis

Statistical analyses were performed using GraphPad program (Abacus Concepts). Results were expressed as mean ± SEM or SD; a *P* value ≤ 0.05 was indicative of statistical significance. The nonparametric Mann-Whitney *U*-test, the Wilcoxon matched-pairs signed-rank test and paired Student’s *t*-test were used. We used two-tailed test for all analyses.

## Data availability

The sequencing data Amplicon-Seq can be found in the European Nucleotide Archive (ENA) database with accession number SUB9171641.

## Acknowledgments

We thank Kabuki subjects and their families that made this study possible with their participation; we thank M.R. Montagna, S. De Simone (Institute of Endotypes in Oncology, Metabolism and Immunology (IEOMI), National Research Council of Italy) and Antonio Amendola (Department of Molecular Medicine and Medical Biotechnology University of Naples “Federico II”) for their technical and experimental support.

This work was supported by grants from Ministry of Education, University and Research (MIUR) PRIN 2022KT2HBJ, PRIN-PNRR 2022C5KBT, European Union - Next Generation EU “PE8 Ageing Well in an ageing society – AGE-IT” Investment 1.3 (Partenariato Esteso - PE0000015) to VDR; Italian Ministry of Health, Italian National Recovery and Resilience Plan (NRRP), Project Code: PNRR-MCNT2-2023-12378040, M6/C2_CALL 2023, funded by Ministero della Salute and MUR NRRP Extended Partnership (INF-ACT no. PE00000007) to GM; Ricerca Corrente by the Italian Ministry of Health, Progetto MiCrO_Care - CUP E63C23002420002, finanziato dalla Regione Campania con Fondo di Rotazione ex L. 183/1987 - Avviso Malattie Rare to G.Merla. IC acknowledges support by FISM - Fondazione Italiana Sclerosi Multipla cod.2020/BC/001 and financed or co-financed with the “5 per mille” public funding. APezone acknowledges support by the University Research Funding Program (FRA) 2022 of the University of Naples Federico II”.

## Author Contributions

**Conceptualization:** V.D.R., I.C. and A.Pe. **Data curation:** A.C., A.L., C.F., A.Po., S.B., F.P., F.G., B.D.S., A.R., M.N.G., A.Pe. and V.D.R. **Formal analysis:** A.C., A.L., C.F., V.M., A.Po., A.L.F., D.Ma, A.S., F.P., A.Pe. and V.D.R. **Funding acquisition:** V.D.R., A.Pe., G.Ma and G.Me. **Investigation:** A.C., A.L., V.M., C.F., A.Po., S.B., F.P., F.G., G.G., E.D., G.T.M., G.S., M.D.M., C.P., D.M., A.R., M.N.G., G.Ma., R.I., G.Me., A.Pe. and V.D.R. **Methodology:** A.C., A.L., C.F., A.Po., F.P., G.Ma., G.Me., A.Pe., and V.D.R. **Project administration:** V.D.R. and A.Pe. **Resources:** A.Po., F.P., G.T.M., G.S., M.D.M., C.P., D.M., G.Ma., R.I., G.Me., A.Pe., and V.D.R. **Software:** A.Pe. **Supervision:** A.Po., A.Pe., and V.D.R. **Validation:** A.C., A.L., V.M., C.F., S.B., F.G., R.B., I.C., A.Pe., and V.D.R. **Visualization:** A.C., A.L., A.Pe. and V.D.R. **Writing-original draft:** A.C., A.L., R.B., A.Pe., I.C., and V.D.R. **Writing-review and editing:** A.C., A.L., G.V., G.Spa., R.B., I.C., A.Pe., and V.D.R.

## Expanded View Figure Legends

**Expanded View Figure 1.**
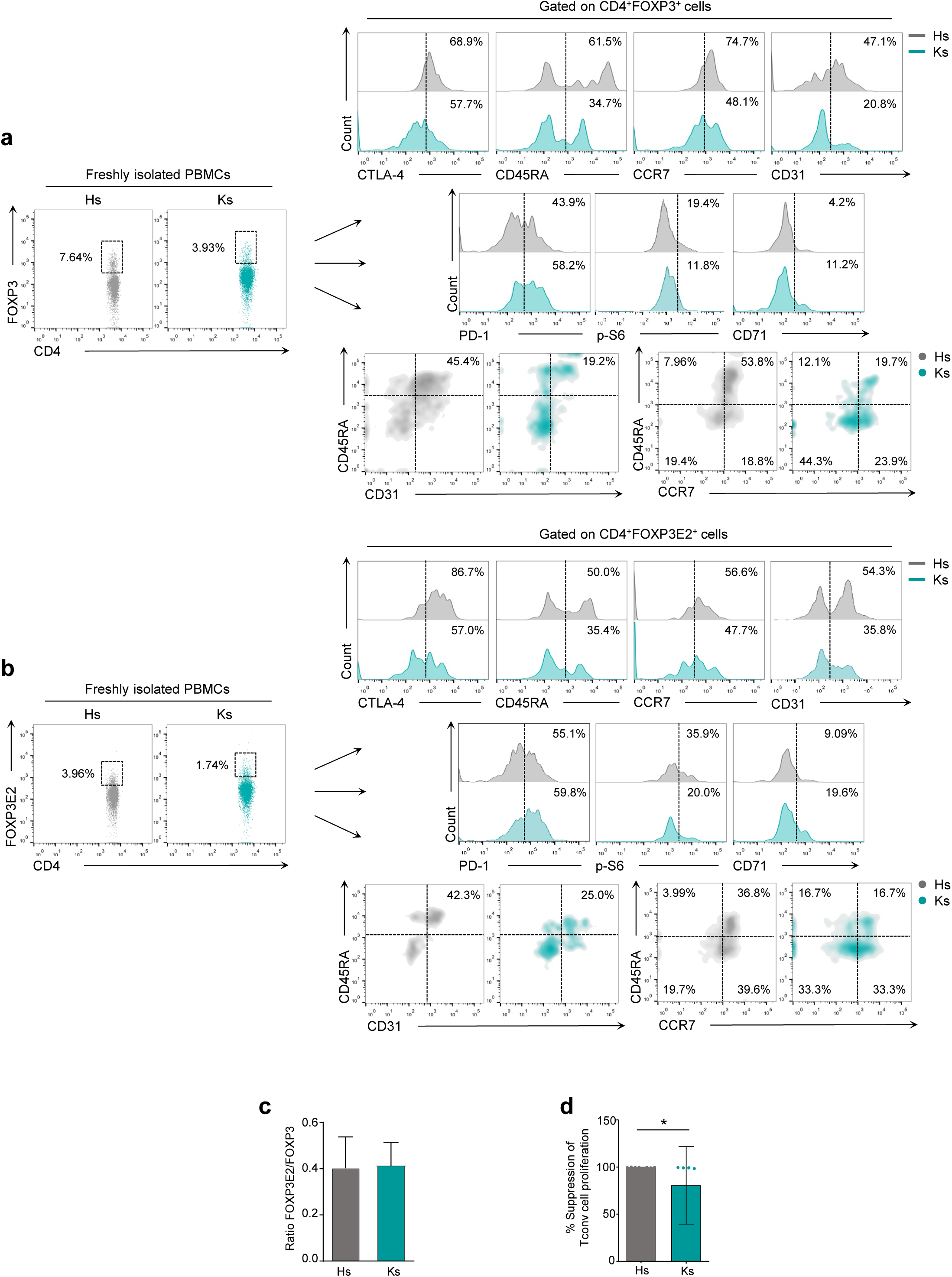
Treg cell frequency, expression of Treg-cell lineage markers and suppressive function in Ks compared to Hs. Flow cytometry analyzing the expression of (**a**, upper left panel) CD4^+^FOXP3^+^ Tregs, (**b**, lower left panel) CD4^+^FOXP3E2^+^ Tregs, and (**a**,**b**, upper right panel) Treg-cell lineage markers in freshly isolated PBMCs from Hs (grey dots and lines) and Ks (teal dots and lines).(**c**) FOXP3E2/FOXP3 ratio evaluated as the ratio between CD4^+^FOXP3E2^+^ Tregs to CD4^+^FOXP3^+^ Tregs in Hs (grey column) and Ks (teal column). Numbers in the plots indicate percentage (%) of positive cells; one representative experiment out of *n*=20 and *n*=13 independent experiments for Hs and Ks, respectively, from at least 8 Hs and 5 Ks. (**d**) % Suppression of Tconv cell proliferation calculated as: (cpm of Tconvs alone – cpm of Tconvs treated with Tregs)/cpm of Tconv cells alone) *100. Tconvs (alone or co-cultured with Tregs) were TCR-stimulated with anti-CD3/anti-CD28 mAbs (0.5 bead per cell) for 60 hours and plated at a 1:1 Tconv:Treg ratio. Data are from n=8 and n=5 independent experiments for Hs (grey column) and Ks (teal column), respectively. Each symbol represents an individual data point. Statistical analysis was performed by using Mann-Whitney *U*-test (two tails) (mean±sd); \**P*≤0.05.

**Expanded View Figure 2.**
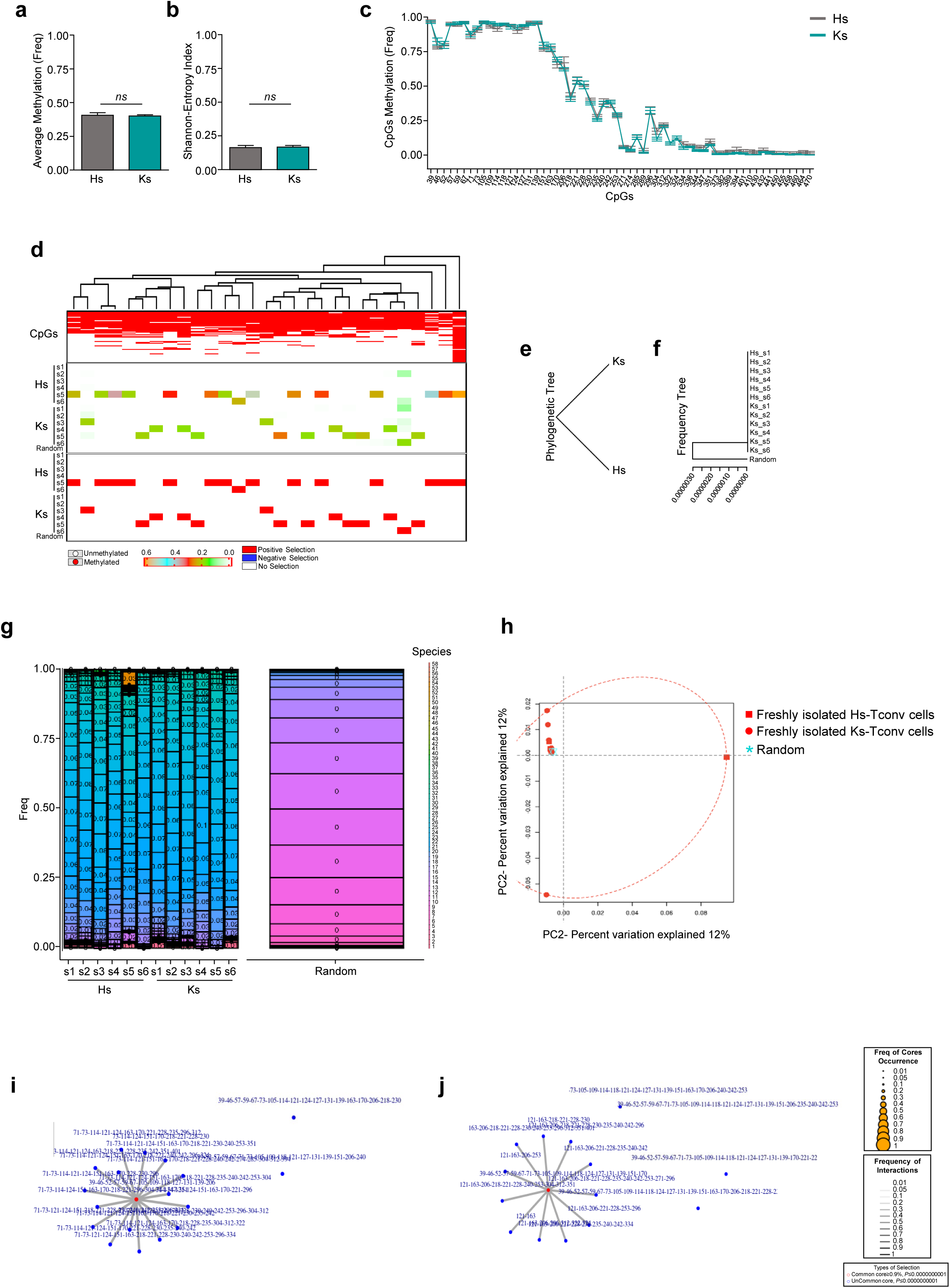
Methylation signature of the *FOXP3* distal enhancer (CNS0) (region I) in Ks and Hs. Bisulfite sequencing of DNA extracted from freshly isolated Tconv cells from Hs and Ks. (**a**) Average methylation, (**b**) Shannon-Entropy Index and (**c**) average methylation of each CpG of *FOXP3* region I in Hs and Ks. (**d**) Structure and frequency of the epialleles from Hs and Ks. The upper section of the panel shows the cluster analysis of all epialleles (methylated, red; unmethylated, white). The central section of the panel shows the frequency of the epialleles and the bottom section displays the epialleles that change significantly (see the color-code legend below the panel). The frequency of epialleles in a Random control is shown far left of each section of the panel (Random). High- or low-frequency or neutral epialleles are shown in red, blue or white, respectively. (**e**) and (**f**) show the Phylogenetic and the Frequency tree of Hs and Ks, respectively. (**g**) Taxonomy of *FOXP3* distal enhancer epialleles (CNS0) (region I) in Hs and Ks (left panel) and the relative Random control (right panel). (**h**) Principal component analysis (PCA) of epialleles derived from Hs and Ks. (**i**) and (**j**) show the complexity of the Hs and Ks-MethCore structures, respectively. The position of CpGs refers to the forward primer. Data are from 6 Hs and 6 Ks. Statistical analysis was performed by using the Student’s *t*-test (mean±SD; **a,b**), (mean±SEM; **c**); *ns*, not significant.

**Expanded View Figure 3.**
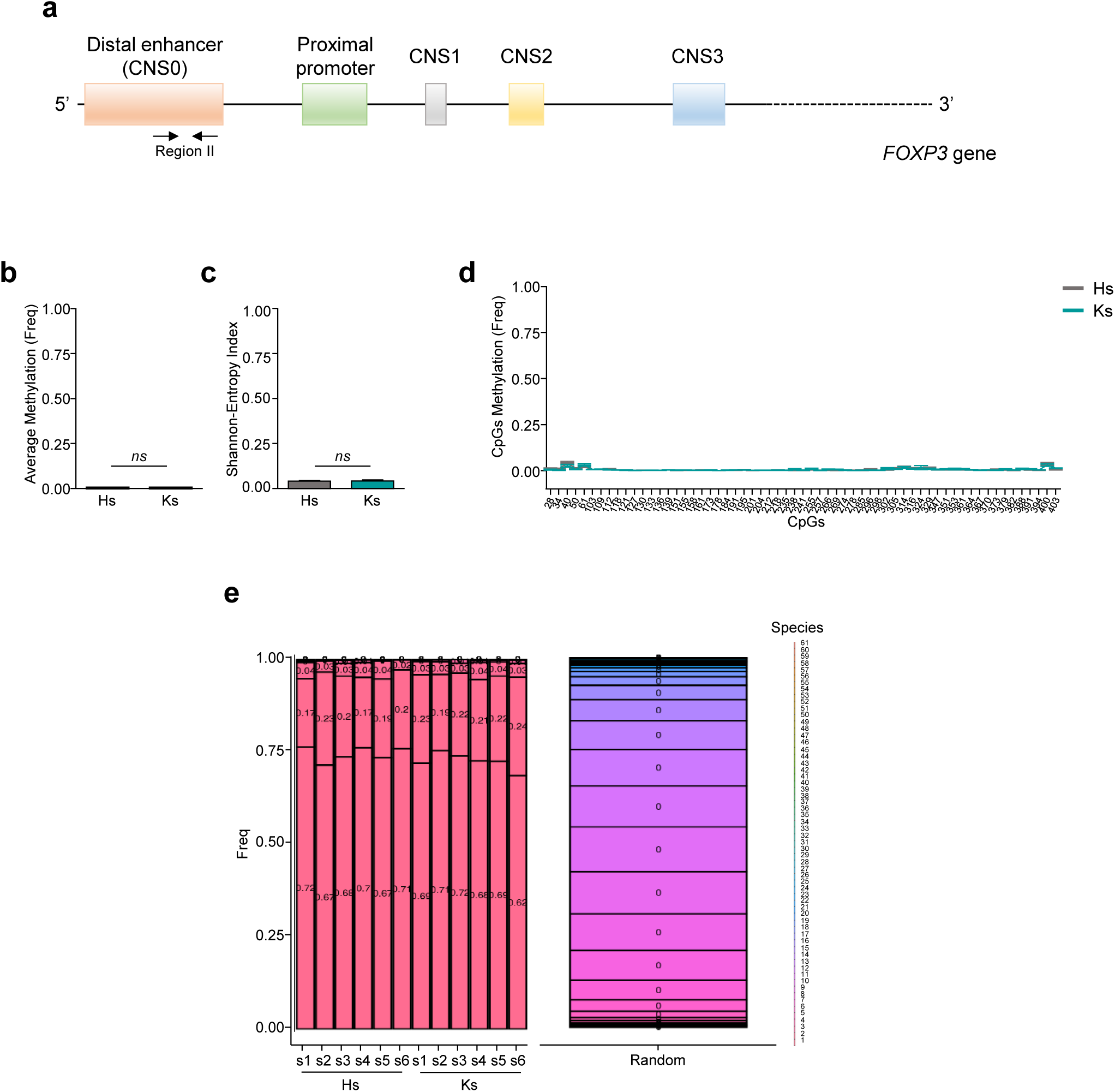
Methylation signature of the *FOXP3* distal enhancer (CNS0) (region II) in Ks and Hs. Bisulfite sequencing of DNA extracted from freshly isolated Tconv cells from Hs and Ks. (**a**) Human *FOXP3* gene structure: black arrows indicate the distal enhancer (CNS0) (region II). Analysis of region II refers to the 61 CpGs present. (**b**) Average methylation, (**c**) Shannon-Entropy Index and (**d**) average methylation of each CpG of *FOXP3* region II in Hs and Ks. (**e**) Taxonomy (left panel) of *FOXP3* distal enhancer epialleles (CNS0) (region II) in Hs and Ks and the relative Random control (right panel). The position of CpGs refers to the forward primer. Data are from 6 Hs and 6 Ks. Statistical analysis was performed by using the Student’s *t*-test (mean±SD; **b**,**c**), (mean±SEM; **d**); *ns*; not significant.

**Expanded View Figure 4.**
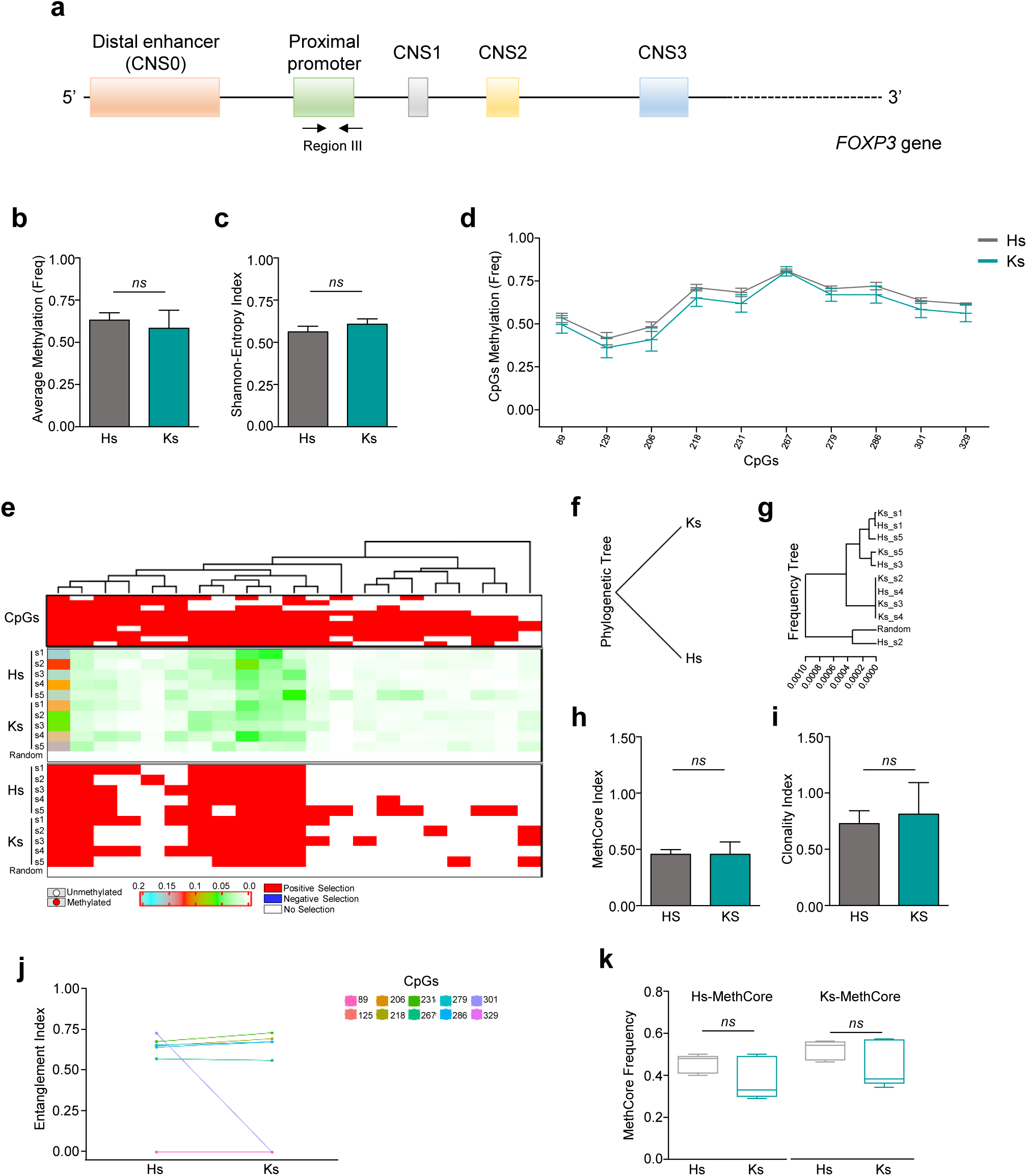
Methylation signature of the *FOXP3* proximal promoter (region III) in Ks and Hs. Bisulfite sequencing of DNA extracted from freshly isolated Tconv cells from Hs and Ks. (**a**) Human *FOXP3* gene structure: black arrows indicate the proximal promoter (region III). Analysis of region III refers to the 10 CpGs present. (**b**) Average methylation, (**c**) Shannon-Entropy Index and (**d**) average methylation of each CpG of *FOXP3* region III in Hs and Ks. (**e**) Structure and frequency of the epialleles. The upper section of the panel shows the cluster analysis of all epialleles (methylated, red; unmethylated, white). The central section panel shows the frequency of the epialleles, and the bottom section displays the epialleles that change significantly (see the color-code legend below the panel). The frequency of epialleles in a Random control is shown far left of each section of the panel (Random). High- or low-frequency or neutral epialleles are shown in red, blue or white, respectively. (**f**) and (**g**) show the Phylogenetic and the Frequency tree of all populations. (**h**) Frequency of Methylated Core (MethCore Index) in the same groups shown in (**b**). (**i**) Frequency of the MethCore normalized to the average methylation (Clonality Index) in each population shown in (**b**). (**j**) Structure of MethCores of Hs and Ks derived from population of significant molecules. Each CpG is labelled with a color code shown in the legend on the right side of the panel (Entanglement Index); the position of CpGs refers to the forward primer. (**k**) Comparison between Hs-MethCore and Ks-MethCore in Hs and Ks, respectively. Data are from 5 Hs and 5 Ks. Statistical analysis was performed by using the Student’s *t*-test (mean±SD; **b**,**c**,**h**,**i**,**k**), (mean±SEM; **d**); *ns*, not significant.

**Expanded View Figure 5.**
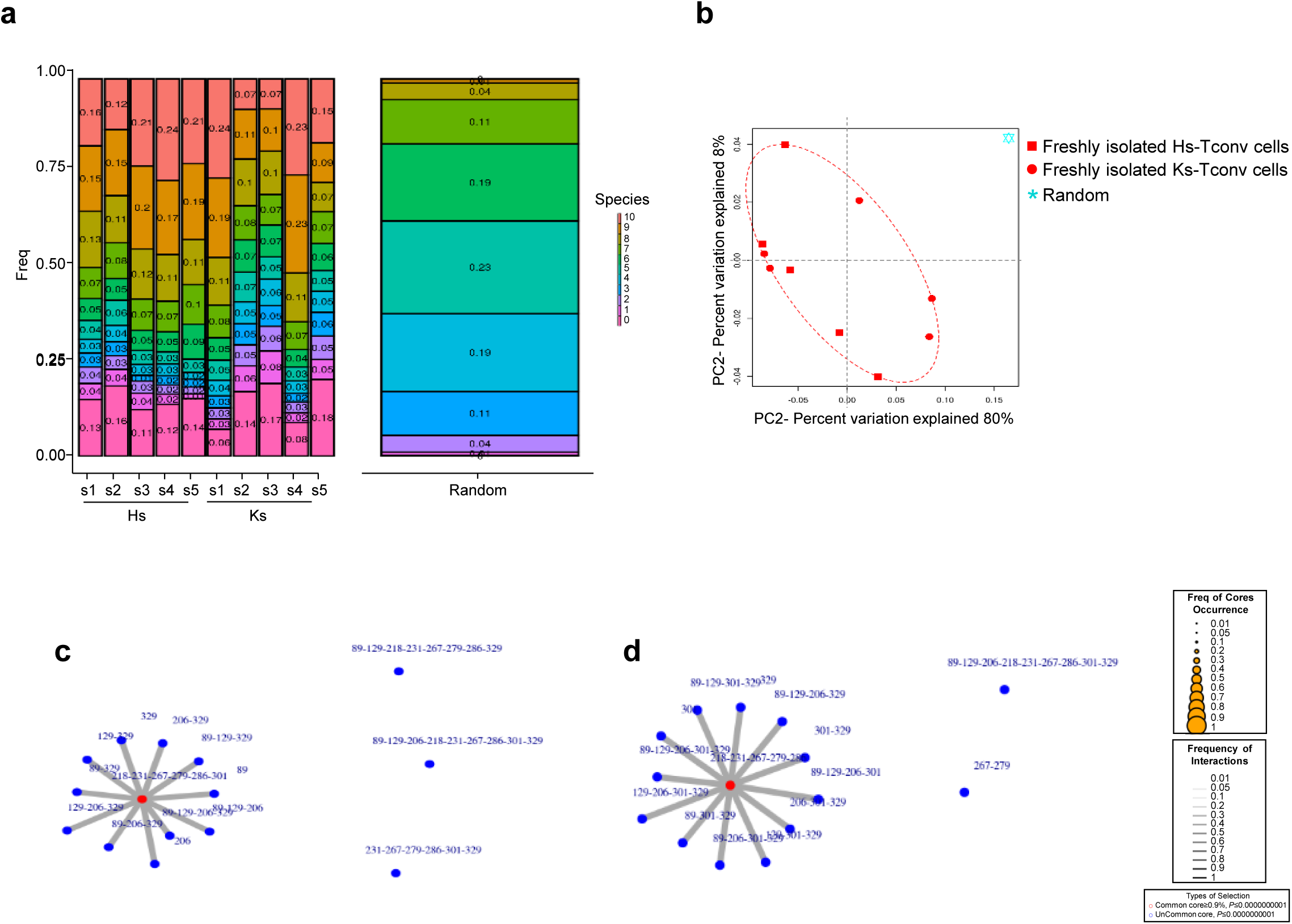
Taxonomy and MethCore structure of the *FOXP3* proximal promoter (region III) in Ks and Hs. (**a**) Taxonomy of *FOXP3* proximal promoter epialleles (region III) in Hs and Ks (left panel) and the relative Random control (right panel). (**b**) Principal component analysis (PCA) of epialleles derived from Hs and Ks. (**c**) and (**d**) show the complexity of Hs- and Ks-MethCore structures, respectively. The position of CpGs refers to the forward primer.

**Expanded View Figure 6.**
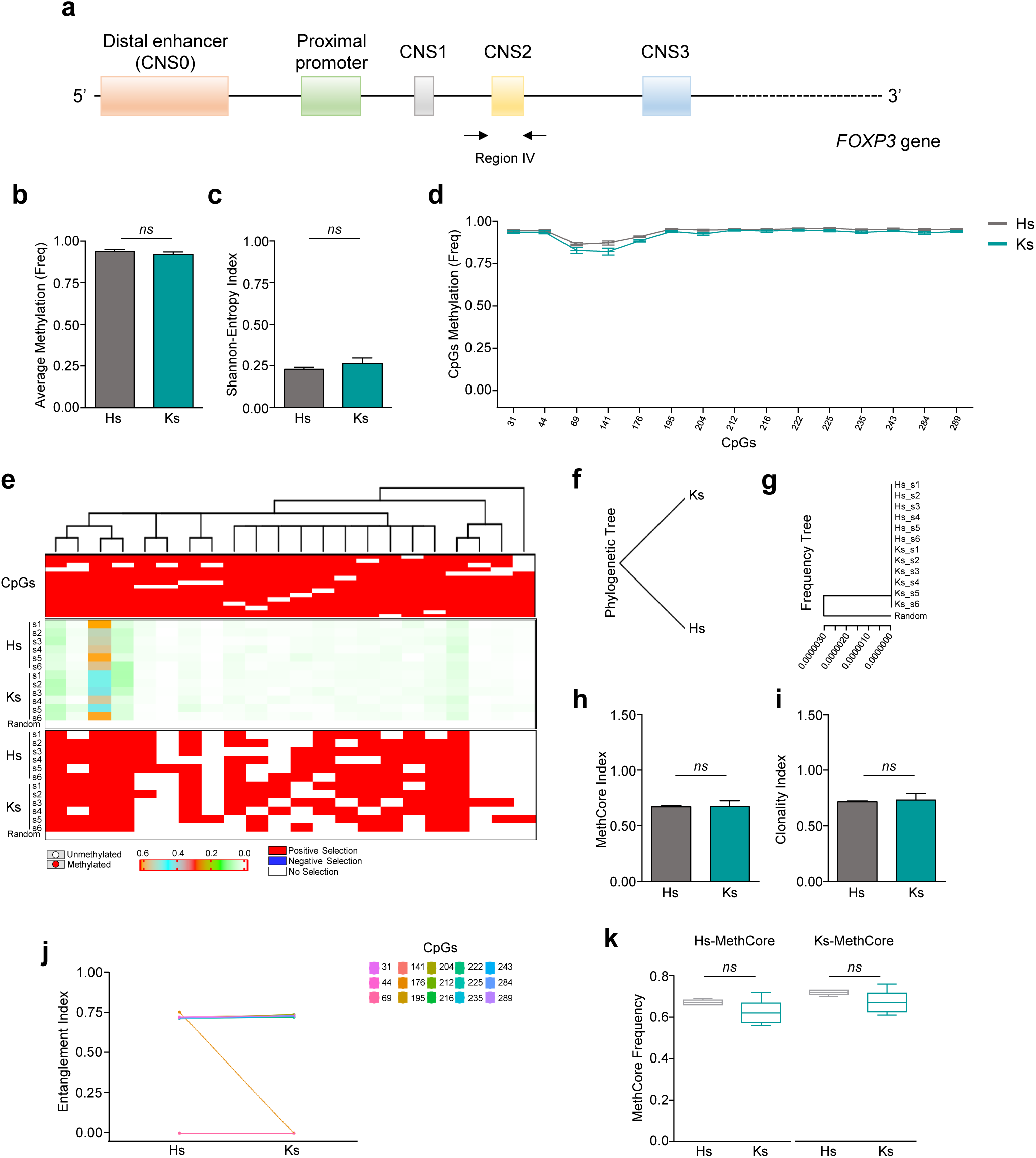
Methylation signature of the *FOXP3* CNS2 (region IV) in Ks and Hs. Bisulfite sequencing of DNA extracted from freshly isolated Tconv cells from Hs and Ks. (**a**) Human *FOXP3* gene structure: black arrows indicate the CNS2 (region IV). Analysis of region IV refers to the 15 CpGs present. (**b**) Average methylation, (**c**) Shannon-Entropy Index and (**d**) average methylation of each CpG of *FOXP3* region IV in Hs (light grey lane) and Ks (teal lane). (**e**) Structure and frequency of the epialleles. The upper section of the panel shows the cluster analysis of all epialleles (methylated, red; unmethylated, white). The central section of the panel displays the frequency of the epialleles, and the bottom section shows the epialleles that change significantly (see the color-code legend below the panel). The frequency of epialleles in a Random control is shown far left of each section of the panel (Random). High- or low-frequency or neutral epialleles are shown in red, blue or white, respectively. (**f**) and (**g**) show the Phylogenetic and the Frequency tree of all populations. (**h**) Frequency of Methylated Core (MethCore Index) in the same groups shown in (**b**). (**i**) Frequency of the MethCore normalized to the average methylation (Clonality Index) in each population shown in (**b**). (**j**) Structure of MethCores of Hs and Ks derived from population of significant molecules. Each CpG is labelled with a color code shown in the legend on the right side of the panel (Entanglement Index); the position of CpGs refers to the forward primer. (**k**) Comparison between Hs-MethCore and Ks-MethCore in Hs and Ks, respectively. Data are from 6 Hs and 6 Ks. Statistical analysis was performed by using the Student’s *t*-test (mean±SD; **b**,**c**,**h**,**i**,**k**), (mean±SEM; **d**); *ns*, not significant.

**Expanded View Figure 7.**
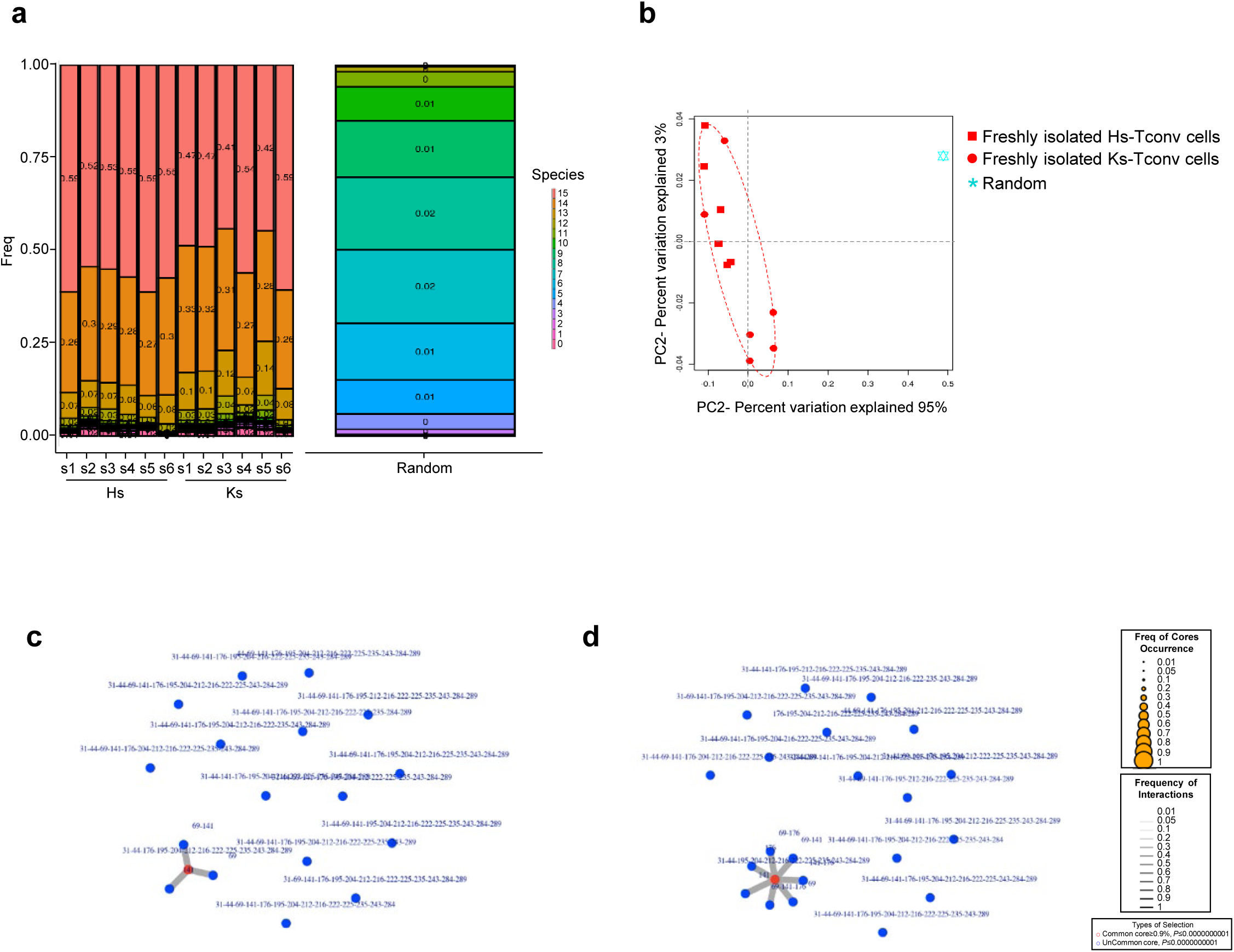
Taxonomy and MethCore structure of the *FOXP3* CNS2 (region IV) in Ks and Hs. (**a**) Taxonomy of *FOXP3* CNS2 epialleles (region IV) in Hs and Ks (left panel) and the relative Random control (right panel). (**b**) Principal component analysis (PCA) of epialleles derived from Hs and Ks. (**c**) and (**d**) show the complexity of Hs- and Ks-MethCore structures, respectively. The position of CpGs refers to the forward primer.

**Expanded View Figure 8.**
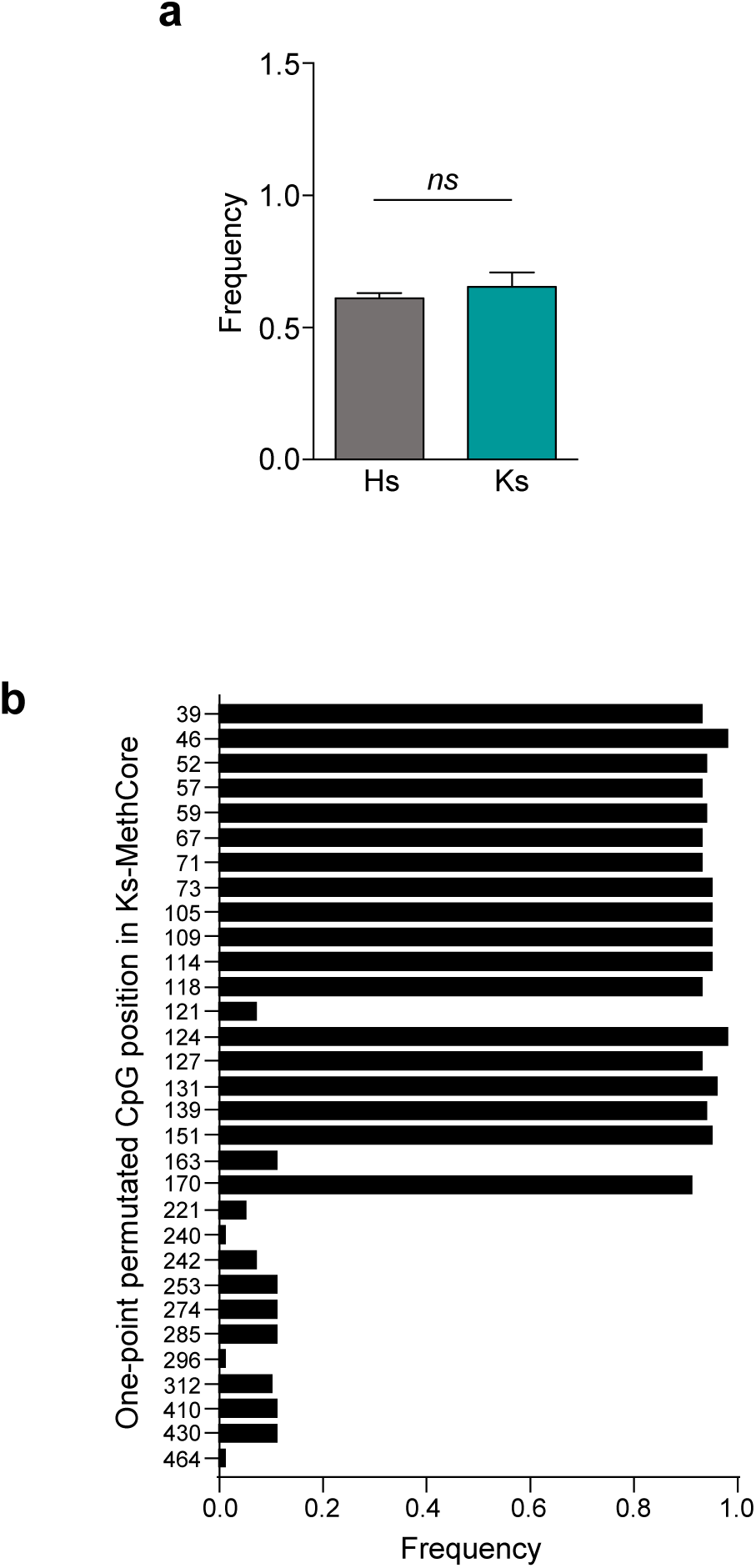
Frequency and stability of epialleles with more than 18 CpGs in Ks and Hs. (**a**) Frequency of methylated classes with more than 18 CpGs at the *FOXP3* distal enhancer (CNS0) (region I) in Ks compared to Hs. (**b**) “One-point permutation test” performed by adding or removing a single CpG in the Ks-MethCore. Among all the 739 possible combinations obtained with the 58 CpGs present in the *FOXP3* distal enhancer (CNS0) (region I), only 151 were significantly more frequent in Ks than in Hs. Analysis of the CpGs making up the 151 combinations shows that 18 CpGs identified as Ks-MethCore have an association frequency of 0.95; *P*<0.05. Statistical analysis was performed using Student’s t-test (mean ± SD); *ns*; not significant.

**Expanded View Figure 9.**
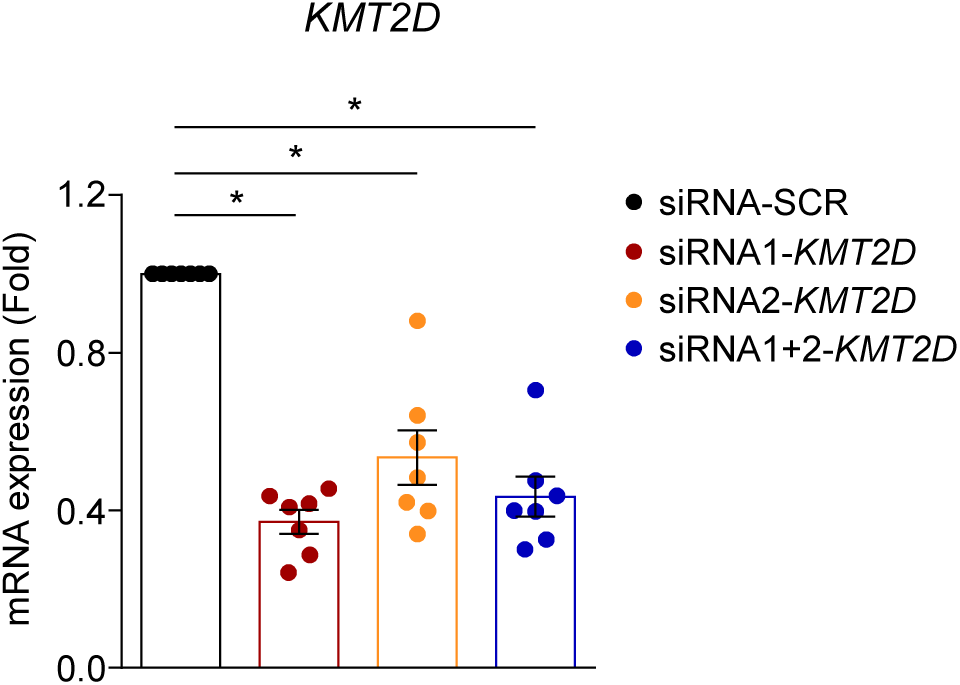
Silencing of *KMT2D* during the generation of inducible (i)Treg cells. Relative mRNA expression of *KMT2D* after its silencing (siRNA1, siRNA2 and both) in Hs-Tconv cells after 24h of TCR stimulation with anti-CD3/anti-CD28 mAbs (0.1 bead per cell). The graph shows *KMT2D* mRNA as fold over siRNA-SCR-Tconv cells. Data are from three independent experiments with technical replicates (*n*=7). Statistical analysis was performed by using Wilcoxon test (two tails) (mean±SEM); \**P*≤0.05.

